# Quorum sensing peptidic inhibitor rescue host immune system eradication: a novel QS infectivity mechanism

**DOI:** 10.1101/2022.10.28.514287

**Authors:** Avishag Yehuda, Einav Malach, Leyla Slamti, Shanny Shuan Kuo, Jonathan Z. Lau, Myung Whan Oh, John Adeoye, Neta Shlezinger, Gee W. Lau, Didier Lereclus, Zvi Hayouka

## Abstract

Subverting the host immune system is a major task for any given pathogen to assure its survival and proliferation. For the opportunistic human pathogen *Bacillus cereus* (Bc), immune evasion enables the establishment of potent infections. In various species of the Bc group, the pleiotropic regulator PlcR and its cognate cell–cell signaling peptide PapR_7_ regulates virulence genes expression in response to fluctuations in population density, *i.e*., a quorum-sensing (QS) system. However, how QS exerts its effects during infections, and whether PlcR confers the immune evading ability remain unclear. Herein, we report how interception of the QS communication in Bc obliterates the ability to control the host immune system. Here we designed a peptide-based QS inhibitor that suppresses PlcR-dependent virulence factor expression and attenuates Bc infectivity in mouse models. We demonstrate that the QS peptidic inhibitor blocks host immune system-mediated eradication by reducing the expression of PlcR-regulated major toxins. Our findings provide the first evidence that Bc infectivity is regulated by QS circuit mediated destruction of the host immunity, thus reveal a new strategy to limit Bc virulence and enhance host defense. This peptidic quorum-quenching agent constitutes readily accessible chemical tool for studying how other pathogen QS systems modulate host immunity and forms a basis for development of anti-infective therapeutics.

## Introduction

Bacterial cell-cell communication or quorum sensing (QS) is a biological mechanism that plays an important role in virulence of numerous bacterial pathogens (Bassler and Losick, 2006; Dunny and Leonard, 1997; Miller and Bassler, 2001). When a critical population mass is reached, detection of signaling molecules allows individual cells to sense the population density and coordinate gene expression of group behaviors, such as antibiotic resistance, competence, sporulation or virulence factor secretion. Gram-negative bacteria predominantly respond to N-acyl homoserine lactones (AHLs), while Gram-positive species mainly rely on cytoplasmic sensors regulated by secreted and re-imported autoinducing short peptides (AIPs; Neiditch et al., 2017; Papenfort and Bassler, 2016; Rajput et al., 2016; Rutherford and Bassler, 2012; Slamti et al., 2014). As targeting such regulators offers the advantage of suppressing bacterial pathogenicity, QS systems have become a powerful candidate for anti-virulence strategies to control infection.

Disruption of QS activity by impairing essential targets in a QS circuit is known as quorum quenching (QQ; Grandclément et al., 2015; Uroz et al., 2009). The AHLs or AIP signals are promising starting point for the development of non-native ligands capable of blocking QS. Therefore, most of QQ chemical strategies comprise direct interference with QS signal molecules, *e*.*g*., by targeting their biosynthesis or receptor interaction (Piewngam et al., 2020; Welsh and Blackwell, 2016). In contrast to antibiotics or antimicrobial agents, which affect the growth of bacteria, QS antagonists could provide an alternative anti-infective therapy with lower selective pressure on the bacterial population to develop resistance (Allen et al., 2014; C., 2000; Rasko and Sperandio, 2010). A wide range of natural or chemically-synthetized molecules have already been presented as QS inhibitors (QSIs) in Gram-negative bacteria (Galloway et al., 2012; Piewngam et al., 2020). More than 100 Gram-negative bacterial species control virulence factor expression using LuxRI-based QS systems (Case et al., 2008). Well-studied examples are LasRI and RhlRI in *Pseudomonas aeruginosa*, with extensive enzymes and small molecule inhibitors that were designed and identified (Mattmann and Blackwell, 2010; Utari et al., 2018). On the other hand, the design of QSIs in Gram-positive bacteria has progressed remarkably only in the last decade (McBrayer et al., 2020). The two-component QS system Agr of *Staphylococcus aureus* was one of the first Gram-positive systems to be targeted (Tal-Gan et al., 2013b, 2013a). To date, an increasing number of studies have shown potent peptide modulators for other species, such as *Streptococcus pneumoniae* (Com system; Yang et al., 2020; Zhu and Lau, 2011), and *Enterococcus faecalis* (Fsr system; (McBrayer et al., 2017, 2018; Nakayama et al., 2013). Recently, we have reported the first synthetic QSIs in *Bacillus cereus* (Bc) that repress the PlcR regulon expression and thus attenuate virulence phenotypes (Yehuda et al., 2018, 2019). Despite the increasing number of studies explored QSIs for their therapeutic potential against Gram-negative and Gram-positive bacterial infections, many of these studies hardly provide the molecular mechanism of QQ.

PlcR is a pleiotropic regulator of virulence genes involved in pathogenicity of the Bc group (Stenfors Arnesen et al., 2008). In addition to the spore-forming Gram-positive bacterial pathogen *sensu stricto*, the Bc group comprises the highly phenotypically and genetically indistinguishable related species: the insect pathogen *Bacillus thuringiensis*, and *Bacillus anthracis*, the causative agent of anthrax (Helgason et al., 2000; Rasko et al., 2005). Bc is also known as a human pathogen responsible for foodborne and opportunistic infections such as endophthalmitis, endocarditis, septicemia, pneumonia, and meningitis, some of which occurring in children and immunocompromised patients (Bottone, 2010; Enosi Tuipulotu et al., 2021; Glasset et al., 2018; Granum and Lund, 1997). These infections are characterized by bacteremia despite the presence of innate immune cells at the site of infected tissue, implying that Bc has developed strategies to overcome the immune response and persist inside its host (Gilois et al., 2007; Hernandez et al., 1998; Tran and Ramarao, 2013; Tran et al., 2011). The precise mechanisms and the bacterial factors allowing Bc to resist to the host immune clearance remain elusive.

Bc utilizes QS to establish opportunistic infection by expressing PlcR-controlled genes, whose functions are related to nutrient acquisition, cell protection, environment-sensing and virulence factors, such as phospholipases, proteases, pore-forming hemolysins, and enterotoxins (Enosi Tuipulotu et al., 2021; Gohar et al., 2008; Ramarao and Sanchis, 2013; Slamti et al., 2014b). The importance of PlcR in regulating Bc pathogenesis and specifically, the toxicity of secreted virulence proteins, was shown by the deletion of the *plcR* gene that reduced the haemolytic activity of Bc, resulting in the loss of its pathogenicity in insect and mouse models of infection (Salamitou et al., 2000). Other studies have linked this to the de-repression of the PlcR-dependent genes, namely, hemolysin BL (HBL), non-hemolytic enterotoxin (NHE), phospholipase C (PC-PLC) and sphingomyelinase (SMase), the latter two being phosphatidylcholine esterases that form cereolysin AB (Beecher and Wong, 2000; Beecher et al., 2000; Enosi Tuipulotu et al., 2021; Pomerantsev et al., 2003; Salamitou et al., 2000). Recent studies highlighted the importance of HBL for Bc pathogenesis (Sastalla et al., 2013), and determined its requirement for mammalian surface receptors LITAF (lipopolysaccharide (LPS)-induced tumor necrosis factor (TNF)-α factor) and CDIP1 (cell death-inducing P53 target 1) to induce cytotoxicity and pyroptosis (Liu et al., 2020).

PlcR activity depends on the binding of the signaling C-terminal heptapeptide PapR_7_ (ADLPFEF) at the end of the exponential growth stage (Slamti and Lereclus, 2002). PapR_7_ is imported by the oligopeptide permease Opp system (OppABCDF; Gominet et al., 2001), binds the tetratricopeptide repeat (TPR)-type regulatory domain of PlcR (Grenha et al., 2013), and promotes recognition of the PlcR box, resulting in transcriptional activation of the target genes. Regulation of the PlcR – PapR system is quite complex; When PlcR is activated, it triggers a positive feedback loop that up-regulates its own transcription and that of genes under its control, including *papR* (Agaisse et al., 1999; Gohar et al., 2002, 2008; Lereclus et al., 1996; Slamti and Lereclus, 2002).

We recently studied the PlcR – PapR activation in Bc at the molecular level (Yehuda et al., 2018). This work revealed the first set of synthetic 7-mer PapR-derived peptides with the potential to serve as QQ agents of Bc QS system. Inhibition of the PlcR regulon activity abrogated the expression of virulence factors, as reflected by a loss of the hemolytic activity without affecting bacterial growth. Next, through multiple amino acids substitution screening, we discovered the first active inhibitor PapR_7_ – dE_6_, with one substitution of D-amino acid instead of glutamic acid (E) at position six of the C-terminus of PapR_7_ (ADLPFEF), indicating that the D-Glutamic substitution was essential for designing potent PlcR antagonist (Yehuda et al., 2019). Many questions remain about the mode of action of the peptide-based PlcR modulators that we have developed, and their potential utility in attenuating virulence during host infection. Additionally, little is known about the molecular mechanisms underlying Bc pathophysiology during host infection. Therefore, in the current study we aimed to use our chemical PlcR inhibitors as valuable tools to investigate the role of QS and the pleiotropic regulator PlcR in colonization and infection by Bc.

In this study, we attempted to optimize the potency of the inhibitory peptide PapR_7_ – dE_6_, through substitution of different amino acids at the first position (Ala1), in addition to D-Glutamic acid at position six of PapR_7_. Through this screening, we were able to develop a new potent PlcR inhibitor termed PapR_7_ – A1M:dE_6,_ containing a methionine replacement of Ala1 position. Our results show that PapR_7_ – A1M:dE_6_ effectively inhibits the expression of PlcR-regulated virulence factors *in vitro* with an IC_50_ value of 200 nM and attenuates Bc infectivity in three mouse models of Bc infection. By applying the new inhibitor to investigate the molecular mechanism underlying its therapeutic effect, we revealed new insights on the interactions between Bc and host macrophages and neutrophils. Moreover, we have identified the bacterial factors allowing Bc to resist these immune cells, thus, providing the first evidence implicating bacterial QS system in impairing the immune system activity during infection.

## Materials and methods

### 1. Bacterial strains and growth conditions

Bacterial strains used in this study: *B. thuringiensis* 407 Cry^−^ *plcA’Z* (Bt A’Z) and the PapR null-mutant 407 Cry^−^ Δ*papR plcA’Z* (Bt ΔpapR A’Z) strains, containing a transcriptional fusion between the promoter of *plcA* and the *lacZ* reporter gene, as described previously (Lereclus et al., 1989; Slamti and Lereclus, 2002). The *B. thuringiensis* 407 Cry^−^ Δ*plcR* and Δ*plcRpapR* (Bt ΔplcR; ΔplcRpapR) mutant strains (Bouillaut et al., 2008; Salamitou et al., 2000). *B. cereus* strain ATCC 14579 (Bc; Ivanova et al., 2003). Unless otherwise noted, all bacterial strains were grown in a modified LB medium (16 g/L tryptone, 8 g/L yeast extract, 5 g/L NaCl) at 37°C and stored at −80°C in LB containing 25% glycerol. Kanamycin (200 μg/mL) was used for the selection of Bt strains.

### 2. Solid phase peptide synthesis methodology (SPPS)

Solid-phase peptide synthesis (SPPS) was performed as previously described (Yehuda et al., 2018, 2019). Briefly, all peptides were synthesized in SPE polypropylene single-Fritted tubes (Altech), using standard Fmoc-based SPPS procedures on Rink Amide resin (substitution 0.6 mmol/g, 250 μmol). De-protecting of Fmoc group was conducted with 20% (v/v) piperidine diluted in dimethylformamide (DMF) followed by heating to 80°C in a microwave (MARS, CEM, United States; 2-min ramp to 80°C, 4-min hold at 80°C). To couple each amino-acid, Fmoc-protected amino acids (4 equiv. relative to the overall loading of the resin), were dissolved in DMF and mixed with 2-(1H-benzotriazol-1-yl)-1,1,3,3-tetramethyluronium hexafluorophosphate (HBTU; 4 equiv.) and diisopropylethylamine (DIEA; 4 equiv.). The solution was allowed to pre-activate for 5 min before being added to the resin and heated to 70°C in a multimode microwave (2-min ramp to 70°C, 4-min hold at 70°C). After each coupling/deprotection step, the resin was drained and washed with DMF (3 × 5 mL). Upon synthesis completion, the peptides were cleaved from the resin [(95% trifluoroacetic acid (TFA), 2.5% triisopropylsilane (TIPS), 2.5% deionized water], re-suspended in 20% acetonitrile [(ACN) in water (v/v)], and lyophilized before high-performance liquid chromatography (HPLC) purification.

### 3. Peptide purification

Crude peptides were purified using Reverse-Phase (RP)-HPLC. The crude peptides were dissolved in 20% ACN in water (v/v) or dimethyl sulfoxide (DMSO). For preparative and analytical RP-HPLC work, a semi-preparative Phenomenex Kinetex C18 column (5 μm, 10 × 250 mm) or analytical Phenomenex Gemini C18 column (5 μm, 4.6 mm × 250 mm, 110 Å) was used, respectively. Standard RP-HPLC conditions were as follows: flow rates = 5 mL min^−1^ for semi-preparative separations and 1 mL min^−1^ for analytical separations; mobile phase A = 18 MΩ water + 0.1% TFA; mobile phase B = ACN. Purities were determined by integration of peaks with UV detection at 220 nm using a linear gradient (first prep 5% B → 65% B over 60 min and second prep 26% B → 36% B over 20 min). The purity of the tested peptides was determined using a linear gradient (5% B → 65% B over 60 min). Only peptide fractions that were purified to homogeneity (>95%) were used for the biological assays. MALDI-TOF spectrometry (Bruker Daltonik, Germany) was used to validate the synthesized peptides molecular weight. The purified peptides were lyophilized and stored at -20°C until use.

### 4. Analysis of PlcR regulon expression using β-galactosidase assay

#### 4.1. PlcR activation studies

Bt ΔpapR A’Z cells were grown overnight in LB medium with selective antibiotic. The cells were diluted 10^−3^ in modified LB to a final volume of 1 L and incubated at 37°C with shaking (200 rpm) until onset of the stationary phase of bacterial growth (OD_600_ 3 ± 0.5). 10 μM of each synthetic A1-substituted:dE_6_ peptides were added to 2 mL aliquots of culture, and incubated for 1 h before centrifugation (Eppendorf centrifuge R5810, 4000 rpm for 5 min), followed by quantification of β-galactosidase activity.

#### 4.2. PlcR inhibition studies

Bt A’Z cells were grown overnight in LB medium. The cells were diluted 10^−3^ in modified LB to a final volume of 1 L and incubated at 37°C with shaking (200 rpm) until the late-exponential of bacterial growth (OD_600_ 1.8 ± 0.1). Different concentrations of synthetic peptides were added to 2 mL aliquots of culture, incubated (1 h) and centrifuged (Eppendorf centrifuge R5810, 4000 rpm for 5 min) for β-galactosidase activity quantification. GraphPad Prism 8 was used to calculate the IC_50_ values, which are the concentration of an inhibitor necessary for 50% inhibition.

#### 4.3. β-galactosidase assay

β-galactosidase activity was measured as described previously (Yang et al., 2017), with a few modifications. Briefly, 200 μL aliquots from 2 mL treated cultures were added in triplicate to a clear 96-well microtiter plate, and OD_600_ was measured (Tecan infinite Pro Plate reader). To quantity the β-galactosidase activity, cells were lysed by incubating for 30 min at 37°C with 20 μL of 0.1% Triton X-100. In a new plate, 100 μL of Z-buffer solution (60 mM Na_2_HPO_4_, 40 mM NaH_2_PO_4_, 10 mM KCl, and 1 mM MgSO_4_, pH 7) containing 2-nitrophenylbeta-D-galactopyranoside (ONPG; at a final concentration of 0.4 mg/mL) was added, followed by 100 μL of lysate, and the plate was incubated for maximum 1 h at 37°C. The reaction was stopped by adding 50 μL of 1 M sodium carbonate solution (Na_2_CO_3_), and the OD 420_nm_ and OD 550_nm_ measured. Each experiment was carried out at least three times, with three replicates per treatment in each experiment. Miller units were calculated using the following formula:

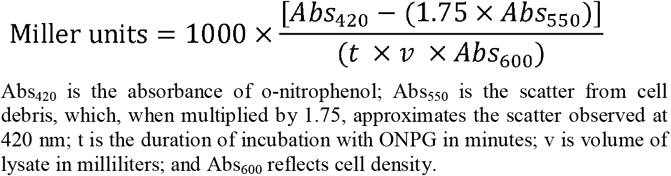

In the Bt ΔpapR A’Z strain, the *plcA* promoter activity was very low and considered as a baseline. In Bt A’Z strain, the untreated bacteria were considered as 100% of activation and the results were normalized accordingly.

### 5. Hemolytic assay on sheep blood agar plates

Haemolytic activity of Bc was examined on trypticase soy agar plates containing 5% washed sheep red blood cells (Hy-Labs, Rehovot, Israel). Overnight cultures were diluted 10^−3^ in modified LB to a final volume of 1 L and incubated at 37°C with shaking (200 rpm) until the end of the lag phase of bacterial growth (OD_600_ 0.1 ± 0.03). Different concentrations of synthetic peptides were added to 2 mL aliquots of bacterial culture and incubated for 2.5 h. The cultures were then centrifuged (Eppendorf centrifuge R5810, 4000 rpm for 5 min), and the supernatants were collected and filtered (0.2 μm filter). 20 μL of filtered supernatants were placed into holes in sheep blood agar plate and incubated for 24 h at 37°C.

### 6. Cell cultures

#### 6.1. Macrophages cell line

RAW 264.7 mouse macrophages were kindly provided by Dr. Sharon Schlesinger. Cells were cultured in Dulbecco’s modified Eagle’s medium (DMEM, Sigma) supplemented with 10% (v/v) fetal bovine serum (FBS), 1% (v/v) Penicillin-streptomycin and 1% (v/v) L-glutamine (Biological Industries) at 37°C in a 5% CO_2_ humidified incubator. The day prior to use, cells were detached by gentle scraping, counted with a haemocytometer to a concentration of 5 × 10^5 cells/mL and seeded into 96-well microplate (Nunc, Thermo Scientific).

#### 6.2. Isolation of bone marrow-derived mouse neutrophils

Bone marrow-derived neutrophils were isolated from C57BL/6 mice (7 week old, males, n=6) as previously described (McGill et al., 2021; Swamydas and Lionakis, 2013). Briefly, mice were euthanized according to approved procedure by the Hebrew University of Jerusalem Institutional Animal Care and Use Committee (IACUC). Euthanized mice were disinfected and dissected to isolate the femur. Muscles on the femur were gently removed with scalpel and scissors while avoiding damage to the head of the femur. The bones were rinsed in 70% ethanol and washed with cold Dulbecco’s phosphate-buffered saline (DPBS, Biological Industries). The epiphyses of the long bones were cut, and bone marrow cells were isolated by scraping the inner lining of the shaft and then flushing the shaft with PBS using a 25-gauge needle into a 50 mL tube fitted with a 100 μm filter. Bone marrow cells were pelleted by centrifuging at 1500 rpm for 5 min (Eppendorf centrifuge R5810). Red blood cells were lysed by resuspending the cell pellet in 1 mL of red blood cell lysis solution (150mM NH_4_Cl, 10mM NH_4_HCO_3_, 2nM EDTA) for 5 min at room temperature and then neutralized with 9 mL PBS. Cells were centrifuged and resuspended in RPMI-1640 (Sigma) supplemented with 10% FBS. The yield was determined by counting the cells with a haemocytometer. Lipopolysaccharide (LPS; 2 μg/mL, Thermo Scientific) was added to activate the bone marrow neutrophils.

### 7. Samples preparation for cell-free supernatant cytotoxicity studies

Overnight cultures of Bc or Bt mutant strains (in the presence of antibiotic selection) were diluted 10^−3^ in modified LB to a final volume of 1 L and incubated at 37°C with shaking (200 rpm) until the end of the lag phase of bacterial growth (OD_600_ 0.1 ± 0.03). Different concentrations of synthetic peptides (0.5-10 μM) were added to 2 mL aliquots of bacterial culture and incubated for 2.5 h. For cytotoxicity assessment during the bacterial growth phase, cells were pre-treated with 50 μM of synthetic peptides and incubated for up to 5 h. The cultures were then centrifuged (Eppendorf centrifuge R5810, 4000 rpm for 5 min), and the supernatants were collected and filtered (0.2 μm filter). All supernatants were diluted in phenol-free DMEM or RPMI-1640 (Sigma) to a final concentration of 12.5% (v/v; medium/cell-free supernatant), and incubated with either macrophages or neutrophils for cytotoxicity assessment.

### 8. Cytotoxicity toward immune system cells

#### 8.1. Analysis of macrophage cytotoxicity and viability

RAW 264.7 mouse macrophages (5 × 10^5 cells/mL), were resuspended in phenol-free DMEM and incubated for 24 h with synthetic peptides at two-fold serial dilutions, or with 12.5% (v/v) cell-free supernatant from Bc or Bt mutant strains. For cell-free supernatant cytotoxicity studies, the control macrophages were challenged with equal volumes of modified LB instead of the supernatant. After incubation, media were aspirated and cells viability was determined by addition of 3-(4,5-Dimethylthiazol-2-yl)-2,5-diphenyltetrazolium bromide (MTT, Sigma) at a final concentration of 0.5 mg/mL. Plates were incubated for an additional 90 min, and OD_595_ was measured using a Tecan Infinite Pro plate reader. Cell viability was normalized (expressed as a percentage of the control group, defined as 100%). All experiments were performed in triplicates.

#### 8.2. Analysis of neutrophil cytotoxicity and viability

Bone marrow-derived mouse neutrophils (3 × 10^5 cells/mL RPMI-1640), were incubated for 2 h with synthetic peptides at two-fold serial dilutions, or with 12.5% (v/v) cell-free supernatant from Bc strains. For cell-free supernatant cytotoxicity studies, control cells were challenged with equal volumes of modified LB instead of the supernatant. After incubation, neutrophils were stained with fluorescence-labelled FITC anti-mouse-Ly6G (BioLegend) and propidium iodide (PI, BioLegend) according to the manufacturer’s instructions for 20 min at 4°C. Cells viability was analysed using Flow Cytometry (NovoCyte Quanteon Flow Cytometer, Agilent). Neutrophils were identified as Ly6G^+^ cells and dead cells were identified as PI^+^ cells. Ly6G^+^ and PI^+^ cells were identified through gating and analysed for FITC and PI fluorescence intensities, respectively. Cell viability was determined by comparing the proportion of neutrophil population stained by PI to the population unstained by PI. In cell-free supernatant cytotoxicity studies, cell viability was expressed as a percentage of the control group, which was assumed to be 100%. All experiments were performed in triplicates.

### 9. Mouse infection studies

Mouse studies were performed according to the Guide for the Care and Use of Laboratory Animals of the National Institutes of Health. Animal protocols were vetted and approved by the IACUC at the University of Illinois at Urbana-Champaign. CD-1 mice (6–7 weeks old, both males and females) were purchased from Charles River Laboratory, and mice were acclimated for one week before experimentation. All mice were housed in positively ventilated microisolator cages with automatic recirculating water located in a room with laminar, high efficiency particulate filtered air. The animals received autoclaved food, water, and bedding.

#### 9.1. Mouse toxicity studies

CD-1 mice (cohorts of 5, males and females) were intraperitoneally injected daily with sterile saline (control) or PapR_7_ – A1M:dE_6_ (10 mg/kg, in 100 μL) for 7 days. The weights of each mouse were recorded daily, and group averages calculated. Mice were euthanized on the 7^th^ day, and blood and major organs (*e.g*., lung, liver, etc) were collected. Blood chemistry and complete blood count with differential analysis were performed at the University of Illinois at Urbana-Champaign Veterinary Diagnostic Laboratory. Histopathological analysis of various organs (hearts, lungs, livers, kidneys and spleens) are described in Section 9.7.

#### 9.2. Mouse model of acute pneumonia

CD-1 mice (7 week old, males and females, 8 per cohort) were anesthetized by isoflurane, and intranasally inoculated with 1.72 × 10^6 CFU of Bc. Infected mice were intravenously (retro-orbital) treated twice daily with PBS, 10 mg/kg of PapR_7_ – A1M:dE_6_ or dF_5_. At 24-hours post-infection (hpi), mice were euthanized and lungs harvested, homogenized with an Omni Soft Tissue Tip™ Homogenizer (OMNI International, Tulsa, OK, USA) in 1 mL of sterile PBS. Bacterial burden was determined by serial dilution plating onto LB agar.

#### 9.3. Mouse model of neutropenic acute pneumonia

CD-1 mice (7 week old, males and females, 8 per cohort) were rendered neutropenic by intraperitoneally injection with cyclophosphamide (150 mg/kg) on Day-5 to Day-2 before bacterial challenge. On Day 0, mice were intranasally inoculated with 5.5 × 10^5 CFU of Bc. Infected mice were intravenously treated at 2 hpi with PBS, or 10 mg/kg of PapR_7_ – A1M:dE_6_ or dF_5_, respectively. At 5 hpi, since some mice were under severe respiratory distress and moribund, all mice were euthanized and lung harvested and homogenized in 1 mL of sterile PBS. The bacterial burden in the lung homogenates determined by serial dilution plating onto LB agar.

#### 9.4. Mouse model of soft tissue thigh infection

One day before infection, the right thighs of CD-1 mice (7 week old, males and females, 8 per cohort) were depilated. The thighs were intramuscularly injected with 4.1 × 10^6 CFU of Bc using a 30g needle (in 25 μL). At 2 and 8 hpi, mice were intravenously injected with PBS, 10 mg/kg of PapR_7_ – A1M:dE_6_ or dF_5_. At 24 hpi, mice were euthanized and thigh muscle tissues harvested, homogenized in 1 mL of sterile PBS, and the bacterial burden in the tissue homogenates determined by serial dilution plating onto LB agar.

#### 9.5. Mouse model of neutropenic soft tissue thigh infection

Neutropenic mice with depilated right thigh (see Section 9.3) were intramuscularly injected with 6 × 10^6 CFU of Bc. At 4 and 9 hpi, mice were intravenously injected with PBS, 10 mg/kg of PapR_7_ – A1M:dE_6_ or dF_5_. At 24 hpi, mice were euthanized and thigh muscle tissues harvested and homogenized in 1 mL of sterile PBS. Bacterial burden in the tissue homogenates was determined by serial dilution plating onto LB agar.

#### 9.6. Mouse model of septicemia survival study

Bc (7.2 × 10^8 CFU/ml) were pre-incubated for 30 minutes at 25°C with PBS or 1 mg/mL of of PapR_7_ – A1M:dE_6_ or dF_5_. CD-1 mice (7 week old, males and females, 15 per cohort) were anesthetized by isoflurane, and intravenously inoculated with 100 μL (7.2 × 10^7 CFU) of Bc. Infected mice were treated twice daily intravenously with PBS, or 10 mg/kg of PapR_7_ – A1M:dE_6_ or dF_5_, for 3 days. Mouse mortality were monitored for 120 hpi. Various organs were collected from moribund mice or at 120-hpi for histopathological analysis.

#### 9.7. Histopathological analysis

Following euthanasia with CO_2_ from a compressed source, mouse organs (hearts, lungs, livers, kidneys and spleens) were immediately harvested and incubated in 10% neutral buffered formalin for at least 24 hours. Various organs were embedded in paraffin, sectioned to 5 μm thickness, and stained with hematoxylin and eosin, and imaged using an Olympus DP70 light microscope (Central Valley, PA, USA).

### 10. Sample preparation for extracellular proteome analysis by SDS-PAGE and LC-MS/MS

Overnight cultures of Bc were diluted 10^−3^ in modified LB to a final volume of 1 L and incubated at 37°C with shaking (200 rpm) until the end of the lag phase of bacterial growth (OD_600_ 0.1 ± 0.03). 10 μM of synthetic peptides were added to 2 mL aliquots of bacterial culture and incubated for 2.5 h. The cultures were then centrifuged (Eppendorf centrifuge R5810, 4000 rpm for 5 min), and the supernatants were collected and filtered (0.2μm filter). The extracellular proteins were precipitated using 20% trichloroacetic acid (TCA) overnight at 4°C and harvested by centrifugation at 21,000 x g for 30 min at 4°C. The TCA-precipitated exoproteins were washed in ice-cold acetone, centrifuged (21,000 × g for 15 min at 4°C) and dried for further SDS-PAGE and proteome analysis by LC-MS/MS.

#### 10.1. Analysis of extracellular protein content by SDS-PAGE

The secretome samples were fractionated by one-dimensional SDS-PAGE with the Mini-protean® 3 kit (Bio-Rad, USA) using a 12% resolving gel and a 5% stacking gel at 200 V for 45 min. After electrophoresis, the gel was stained with Coomassie Blue G-250 overnight, de-stained and an image of the gel was captured.

#### 10.2. Analysis of proteome analysis by LC-MS/MS

##### 10.2.1. Cell lysates

Precipitated proteins were solubilized in 100 μL of 8M urea, 10 mM DTT, 25 mM Tris-HCl pH 8.0 and incubated for 30 min at 22°C. Iodoacetamide (55 mM) was added, followed by incubation for 30 min (22°C, in the dark). Trypsin digestion was done using sequencing-grade modified Trypsin (0.4 μg/sample; Promega Corp., Madison, WI) overnight at 37°C. The samples were acidified by addition of 0.2% formic acid and desalted on C18 home-made Stage tips. Peptide concentration was determined by absorbance at 280 nm and 0.3 μg of peptides were injected into the mass spectrometer.

##### 10.2.2. NanoLC-MS/MS analysis

MS analysis was performed using a Q Exactive-HF mass spectrometer (Thermo Fisher Scientific, Waltham, MA USA) coupled on-line to a nanoflow UHPLC instrument, Ultimate 3000 Dionex (Thermo Fisher Scientific, Waltham, MA USA). Peptides dissolved in 0.1% formic acid were separated without a trap column over an 80 min acetonitrile gradient run at a flow rate of 0.3 μl/min on a reverse phase 25-cm-long C18 column (75 μm ID, 2 μm, 100Å, Thermo PepMapRSLC). The instrument settings were as described previously (Scheltema et al., 2014). Data were acquired using Xcalibur software (Thermo Scientific). To avoid a carryover, the column was washed with 80% acetonitrile, 0.1% formic acid for 25 min between samples.

##### 10.2.3. MS data analysis

Mass spectra data were processed using the MaxQuant computational platform, version 2.0.1.0. Peak lists were searched against Uniprot FASTA sequence database from Oct. 31, 2021: *B. cereus* containing 6,331 entries. The search included cysteine carbamidomethylation as a fixed modification, N-terminal acetylation and oxidation of methionine as variable modifications and allowed up to two miscleavages. The ‘match-between-runs’ option was used and the required FDR was set to 1% at the peptide and protein level. Relative protein quantification in MaxQuant was performed using the label-free quantification (LFQ) algorithm algorithm (Cox et al., 2014). Statistical analysis (n=3) was performed using the Perseus statistical package, the Perseus computational platform for comprehensive analysis of (prote)omics data.). Only those proteins for which at least 2 valid LFQ values were obtained in at least one sample group were accepted for statistical analysis by t-test (p < 0.05). After application of this filter, a random value was substituted for proteins for which LFQ could not be determined (“Imputation” function of Perseus). The imputed values were in the range of 10% of the median value of all the proteins in the sample and allowed calculation of p-values.

### 11. Statistical Analysis

Unless otherwise noted, the results are presented as the mean ± SEM. One-way analysis ANOVA of variance, followed by Tukey post hoc comparison was used for statistical analysis. The results were considered to be statistically significant if p < 0.01. For survival analyses, a Kaplan–Meier Log Rank Survival Test was performed using the GraphPad Prism version 9.0.2 software

## Results

### PapR_7_ – A1M:dE_6_ quenches the virulence of Bc and attenuates acute lung infection

We have previously reported the first five synthetic peptidic inhibitors of Bc PlcR-PapR QS system, four of which carrying substitutions of alanine or D-amino acid instead of glutamic acid (E) or phenylalanine (F) at positions 6–7 of the C-terminus of heptapeptide PapR (PapR_7_ – E6A/F7A or PapR_7_ – dE_6_/dF_7_ respectively; Yehuda et al., 2018). To improve the inhibitory activity of the designed PapR_7_ derivatives, we have used multiple amino acids substitution strategy and found that one replacement of Glu6 to its D-enantiomer is sufficient to yield a powerful PlcR antagonist, with the lowest IC_50_ value of 0.977 ± 0.04 μM (Yehuda et al., 2019). Since our previous findings indicated that the first N-termini amino acid (Ala1) is dispensable and thus might be replaced to other type of amino acids in order to improve the potency of the inhibitory peptide PapR_7_ – dE_6_ (Slamti and Lereclus, 2002; Yehuda et al., 2018). The initial set of eight A1-substituted:dE_6_ peptides (listed in Table S1) was synthesized and purified. This set included peptides with eight different amino acid substitutions at Ala1 position, in addition to D-Glutamic acid at position 6 of the PapR heptapeptide.

We tested the peptides ability to modulate the activity of PlcR using *B. thuringiensis* 407 Cry^−^ (Bt 407^−^) as a model bacterium for the Bc group. We used the following reporter strains: *B. thuringiensis* 407^−^ *plcA’Z* (Bt A’Z) and PapR-null mutant *B. thuringiensis* 407^−^ Δ*papR plcA’Z* (Bt ΔpapR A’Z) strains containing *plcA* promoter *lacZ* fusions that have been shown high phylogenic similarity with the Bc reference strain ATCC 14579 and previously used to assess quorum-sensing activity and PlcR regulon expression. (Gominet et al., 2001; Slamti and Lereclus, 2002; Yehuda et al., 2018, 2019). Consistent with prior observations of PapR_7_ – dE_6_ inhibitor, none of the new A1-substituted:dE_6_ peptides activated the PlcR regulon (Figure S1A, Table S1) suggesting their inhibitory activity potential. We next examined their inhibitory activity in Bt A’Z reporter strain at higher concentration (10 μM). Five of the amino acid replacements of alanine residue at PapR_7_ – dE_6_ (Ser, Leu, Try, Asp and Met1), were found to significantly inhibit *plcA’Z* strain (Figure 1B). The IC_50_ values of the derived peptides (Figures 1B) revealed two new inhibitory peptides, PapR_7_ – A1Y:dE_6_ and PapR_7_ – A1M:dE_6_, both having lower IC_50_ values compared to their parent reported PapR_7_ – dE_6_ inhibitor. Interestingly, replacement of Ala1 with methionine significantly increased the inhibitory activity by 5-fold and enhanced its antagonist features (Figures 1B, Table S1). Based on these finding, we replaced Ala1 with methionine in all of our previously-reported Bc inhibitory synthetics PapR_7_-derived peptides (PapR_7_ – E6A, F7A and dF_7_), observing that the Met1 substitution had a significant effect on the inhibition of PlcR regulon expression, reflected in IC_50_ values approximately threefold lower than that of their parent peptidic inhibitors (Figure 1C, Table S2). However, PapR_7_ – A1M:dE_6_ exhibited the lowest IC_50_ value, demonstrating that it is not only the most active inhibitor identified in this series of A1-substituted:dE_6_ derivatives, but also, to our knowledge, the most potent peptidic inhibitor of Bc QS system to be reported.

**Figure 1:**
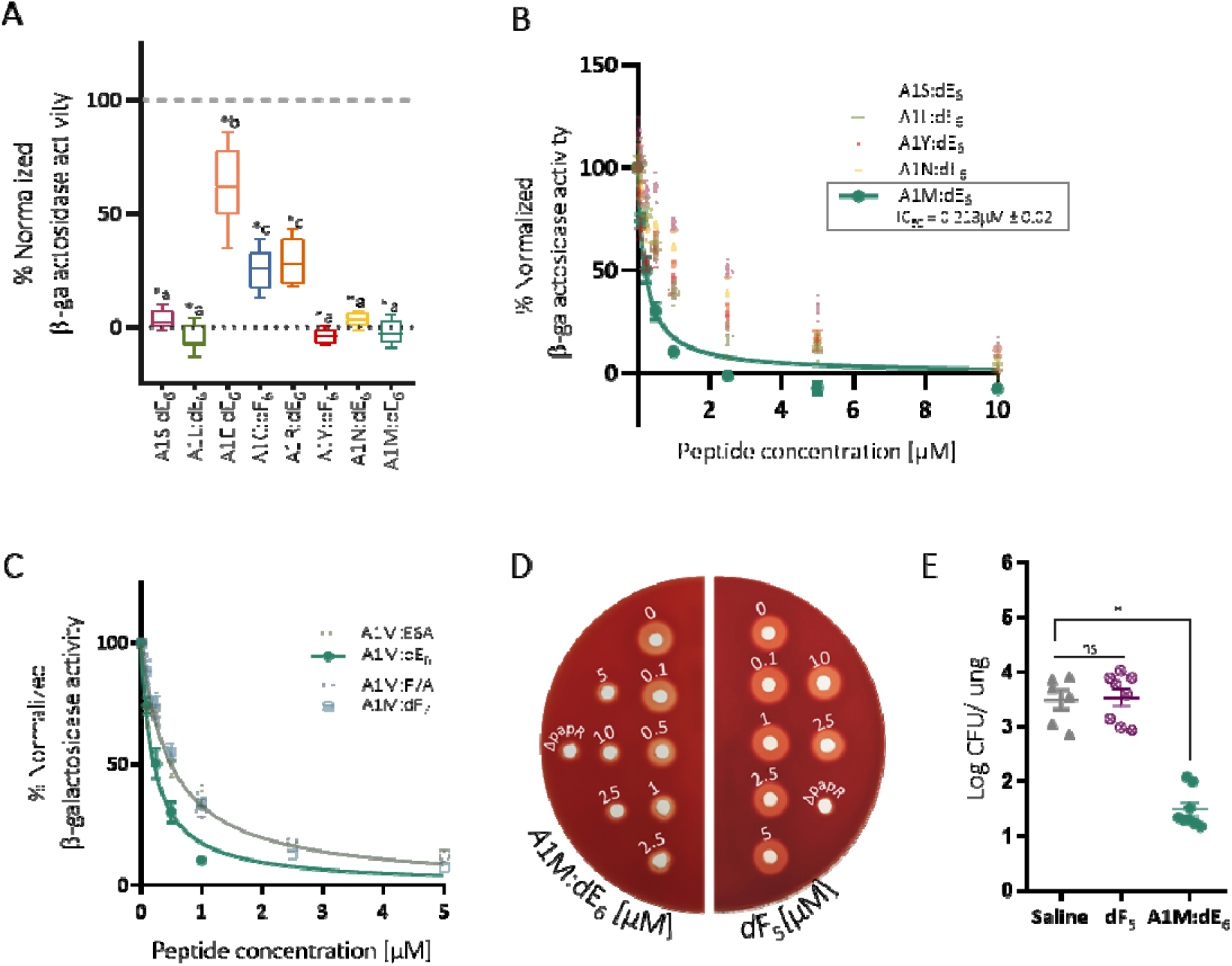
PapR_7_ – A1M:dE_6_ quenches the virulence of Bc and attenuates acute lung infection. (A) β-galactosidase activity of Bt A’Z induced by the addition of 10 μM A1-substituted:dE_6_ peptides, (B) selected strong inhibitory A1-substituted:dE_6_ peptides and (C) A1M-substituted inhibitors in several concentrations normalized to untreated bacterial cells at late-exponential phase of growth (OD_600_ of 1.8 ± 0.1, mean ± SEM, n = 9). *p < 0.01 indicates a statistically significant difference between untreated Bt A’Z versus those treated with A1-substituted:dE_6_ peptides. Different letters indicate statistically significant differences between A1-substituted:dE_6_ peptide treatments (p < 0.01). (D) Representative picture showing the hemolytic activity on sheep blood agar of supernatants from Bc cultures treated with different concentrations of PapR_7_ – A1M:dE_6_ (inhibitory peptide) or PapR_7_–dF_5_ (non-inhibitory peptide). (E) Bacterial loads in the lung of infected mice treated with saline, PapR_7_–dF_5_ and A1M:dE_6_ (mean ± SEM, n = 6-8). *p < 0.01 indicates a statistically significant difference between treatments. ns indicates no statistically significant difference between treatments.

Before evaluating PapR_7_ – A1M:dE_6_ potential utility in attenuating virulence during host infection, we examined its ability to modulate virulence gene expression. Previous studies have shown that Bc pathogenicity is mainly due to synergistic effects of a number of virulence factors that promote intestinal cell destruction and/or resistance to the host immune system. The majority of these exported virulence factors are under the control of PlcR and include several of enterotoxins, phospholipases and hemolysins (Salamitou et al., 2000; Slamti and Lereclus, 2002; Slamti et al., 2004; Visiello et al., 2016). Therefore, we studied the PapR_7_ – A1M:dE_6_ effect on the hemolytic activity of Bc. As shown in Figure 1D; the incubation of of PapR_7_ – A1M:dE_6_ during Bc growth, greatly reduced the hemolysis of sheep red blood cells in a concentration dependent manner. In contrast, the non-inhibitory control peptide, PapR_7_ –dF_5_ (Yehuda et al., 2018), did not affect hemolytic activity profile. These results show the high efficacy of PapR_7_ – A1M:dE_6_ in quenching virulence of Bc *in-vitro*, and might imply on its therapeutic potential during host infection.

Prior to evaluating PapR_7_ – A1M:dE_6_ ability to attenuate Bc infections *in-vivo*, we examined the toxicity of the peptide. CD1 mice (cohorts of five) were intraperitoneally injected daily with PapR_7_ – A1M:dE_6_ (50 mg day^−1^) or saline for one week. Systemic toxicity was assessed by performing blood chemistry and complete blood count (CBC) with differential at the University of Illinois College of Veterinary Medicine Clinical Pathology Laboratory. Their major organs (hearts lungs, livers, spleens and kidneys) were analyzed histopathologically, by using Hematoxylin and Eosin (H&E) staining. Importantly, PapR_7_ – A1M:dE_6_ did not induce any abnormal pro-inflammatory response; with the absence of myelosuppression, renal injury, hepatic toxicity, or other abnormalities (Figure S1 B-C; data not shown). After confirming the safety of PapR_7_ – A1M:dE_6_ treatment, we evaluated its efficacy in a mouse model of acute pneumonia. Mice (8 per cohort) were intranasally inoculated with Bc ATCC 14579 (1.7×10^6^ CFU/mouse). Infected mice were treated twice (2 and 8 hpi) with either saline, or 10 mg/kg of PapR_7_ – dF_5_ (non-inhibitory peptide) and PapR_7_ – A1M:dE_6_, at 24 hpi we evaluated the bacterial burdens in the lungs (Figure 1E). Interestingly, treatment of PapR_7_ – A1M:dE_6_ resulted in a 2.5-logs lower bacterial load than for the saline and PapR_7_ –dF_5_ treated groups (p<0.01). To further, confirm the efficacy of PapR_7_ – A1M:dE_6_ *in-vivo*, the protective effect against Bc was also assessed in mouse model of thigh infection (Figure S1D). Similar to the acute pneumonia model, PapR_7_ – A1M:dE_6_ treated group showed significantly lower bacterial counts in infected thighs, compared to the control treatments. Because PapR_7_ – A1M:dE_6_ is non-bactericidal (Data not shown), the lower bacterial burden in the treated mice suggest that the QSI might have direct effect in preventing Bc to secrete virulence factor that alter the host immune system activity. This might be the reason for the higher microbial load in the control treatment where, the QS secreted virulence factors subvert the immune system activity, resulting in a higher microbial burden.

### PapR_7_ – A1M:dE_6_ protects macrophages against the induced-toxicity of Bc

Next, we wanted to elucidate the molecular mechanism underlying the lower bacterial burdens in the lungs or thighs of mice treated with PapR_7_ – A1M:dE_6_. Because PapR_7_ – A1M:dE_6_ is non-bactericidal, our hypothesis was that the protective activity is mediated by the host immune system, therefore, we first explored the interaction between Bc supernatant with macrophages, focusing on ways in which PapR_7_ – A1M:dE_6_ could prevent the bacterium from overcoming the microbicidal activities of these professional phagocytes. We started by assessing the toxicity of PapR_7_ – A1M:dE_6_, PapR_7_–dF_5_ and PapR_7_ peptides towards macrophages on RAW 264.7 mouse macrophage cell lines using MTT assay. As shown in Figure 2A; incubation of macrophage with up to 400μM of the synthetic peptides for 24 h did not affect the viability of the cells, indicating that at these concentrations, none of the tested peptides was cytotoxic toward macrophage cells.

**Figure 2:**
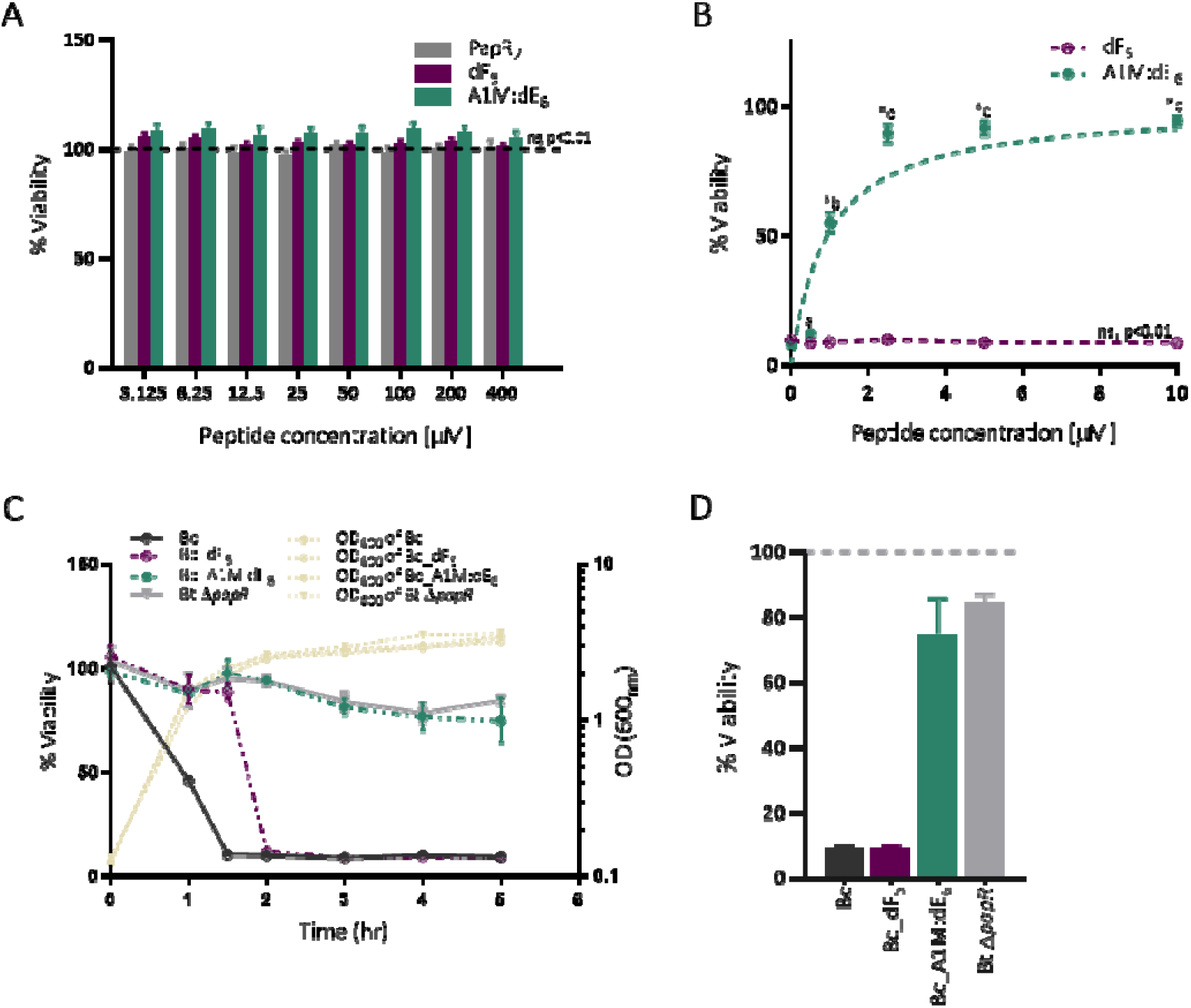
PapR_7_ – A1M:dE_6_ protects macrophages eradication induced by Bc. (A) Cytotoxicity of various peptides (PapR_7_, PapR_7_ – A1M:dE_6_ and dF_5_) on RAW 264.7 mouse macrophages (mean ± SEM, n = 9). ns indicates no statistically significant difference between treatments at different concentration of examined peptides. (B) Macrophage viability after challenge with supernatants from Bc cultures pretreated with PapR_7_ – A1M:dE_6_ and dF_5_ (0.5-10μM, mean ± SEM, n = 9). *p < 0.01 indicates a statistically significant difference between supernatants from untreated Bc cultures versus pretreatment with PapR_7_ – A1M:dE_6_. Different letters indicate statistically significant differences between supernatants of Bc cultures pretreated with different PapR_7_ – A1M:dE_6_ concentrations (p < 0.01). (C) Macrophage viability after challenge with supernatants from Bt ΔpapR A’Z and peptide pretreated Bc cultures (PapR_7_ – A1M:dE_6_ or dF_5_; 50μM) during different bacterial growth phase (data are representative of three independent experiments; mean ± SD, n = 3). (D) Macrophage viability after 5 hours as represented in Panel C (Data are representative of three independent experiments; mean ± SD, n = 3).

Previous studies have shown that cell-free supernatant from Bc cultures is cytotoxic and induced rapid death of macrophages, but the precise regulators and the virulence factors are not fully understood (Ramarao and Lereclus, 2005; Salamitou et al., 2000; Tran et al., 2010, 2011). Therefore, we examined whether supernatants from Bc cultures pre-treated with PapR_7_ – A1M:dE_6_ would exhibit different levels of cytotoxicity towards macrophages. Importantly, addition of the PapR_7_ – A1M:dE_6_ peptide to the growth media was able to restore macrophage viability in a concentration-dependent manner (Figure 2B), compared to untreated Bc culture supernatants or supernatant from BC culture pretreated with the non-inhibitory peptide PapR_7_ –dF_5_, which showed 90% mortality. These results demonstrate that PapR_7_ – A1M:dE_6_ treatment protects macrophages from Bc-mediated toxicity, presumably by preventing the expression and secretion of virulence factors that induce cytotoxicity.

To assess the long-term protective effect of PapR_7_ – A1M:dE_6_, we sampled supernatants from various Bc pre-treated cultures at different time points during bacterial growth, and incubated with macrophage cells. Both native and PapR_7_ – dF_5_ treated supernatants were found to be highly cytotoxic from the point of entry into stationary phase (OD_600_ of ∼ 2.5), and induced rapid cells death (Figure 2C). These findings support our hypothesis that the death of macrophages is related to the secretion QS-dependent virulence factors, which is known to be cell density dependent. However, when the bacteria were pre-treated with PapR_7_ – A1M:dE_6_ the cytotoxicity towards macrophages was abrogated, and the macrophages viabilities were similar to those induced by Bt ΔpapR A’Z mutant strain. Importantly, our results show that the protective effect of PapR7 – A1M:dE_6_ was preserved up to 3.5 hours after entry into the stationary phase of bacterial growth that at that point the secreted virulence factor are at the highest concentration (Figure 2, C-D). To further support the role of PlcR-PapR system in the induction of macrophages, we preformed the experiment with supernatants from Bt ΔplcR and Bt ΔplcRpapR mutant strains (Figure S2, A-B). The supernatants of these mutant strains exhibited very low macrophage killing (less than ∼15%). These results clearly indicate that PlcR-dependent virulence factors secretion plays a major role in Bc cytotoxicity *in-vitro*. Our findings demonstrate the ability of PapR_7_ – A1M:dE_6_ to suppress the toxic effect induced by Bc toward macrophages and thus rescue them from rapid death. We suggest that by preventing the production of PlcR-dependent virulence factors, PapR_7_ – A1M:dE_6_ enables the host immune system to clear Bc during both lung and soft tissue infections.

### PapR_7_ – A1M:dE_6_ confers protective effect on the mice immunity during bacteremia infection

Bc opportunistic infections are characterized by bacteremia (Enosi Tuipulotu et al., 2021; Hernandez et al., 1998; Tran and Ramarao, 2013) hence, we evaluated the efficacy of PapR_7_ – A1M:dE_6_ in reducing mortality in a mouse model of bacteremia caused by the Bc ATCC 14579 strain. CD1 mice (cohorts of 15) were retro-orbitally infected with Bc and treated twice daily with 10 mg/kg of intravenously administered saline, PapR_7_ –dF_5_ or A1M:dE_6_. The bacterial inoculum was pre-incubated for 30 minutes with 1 mg/mL of our examined peptides. For infected mice treated with saline or PapR_7_ –dF_5_, 100% mortality was observed within 10 hours post-infection (Figure 3A). In contrast, mice treated with PapR_7_ – A1M:dE_6_ exhibited significantly higher survival rates even after 5 days and delayed the killing kinetics compared to that of the untreated groups (Figure 3A). Histopathological analysis of hearts tissue showed multiple bacterial microcolonies, associated with tissue damage and mild inflammation in either saline or PapR_7_ – dF_5_ treated mouse (Figure 2, B-C; Figure S3, A-B). Remarkably, a slight to mild degree of neutrophilic alveolitis and hepatitis was observed in the lungs and liver of these cohorts, respectively (Figure S3, C-D). The mild tissue inflammation in the current animal study implied a peracute to acute disease progression, with acute myocardial necrosis and renal tubular necrosis as likely cause of death. In contrast, no overt evidence of intralesional bacterial microcolonies, tissue damage, and inflammation was observed in the Bc infected mice treated with PapR_7_ – A1M:dE_6_ (Figure 2, D-E). Over all, these results highlighting the great therapeutic potential of PapR_7_ – A1M:dE_6_ in attenuating Bc infections.

**Figure 3:**
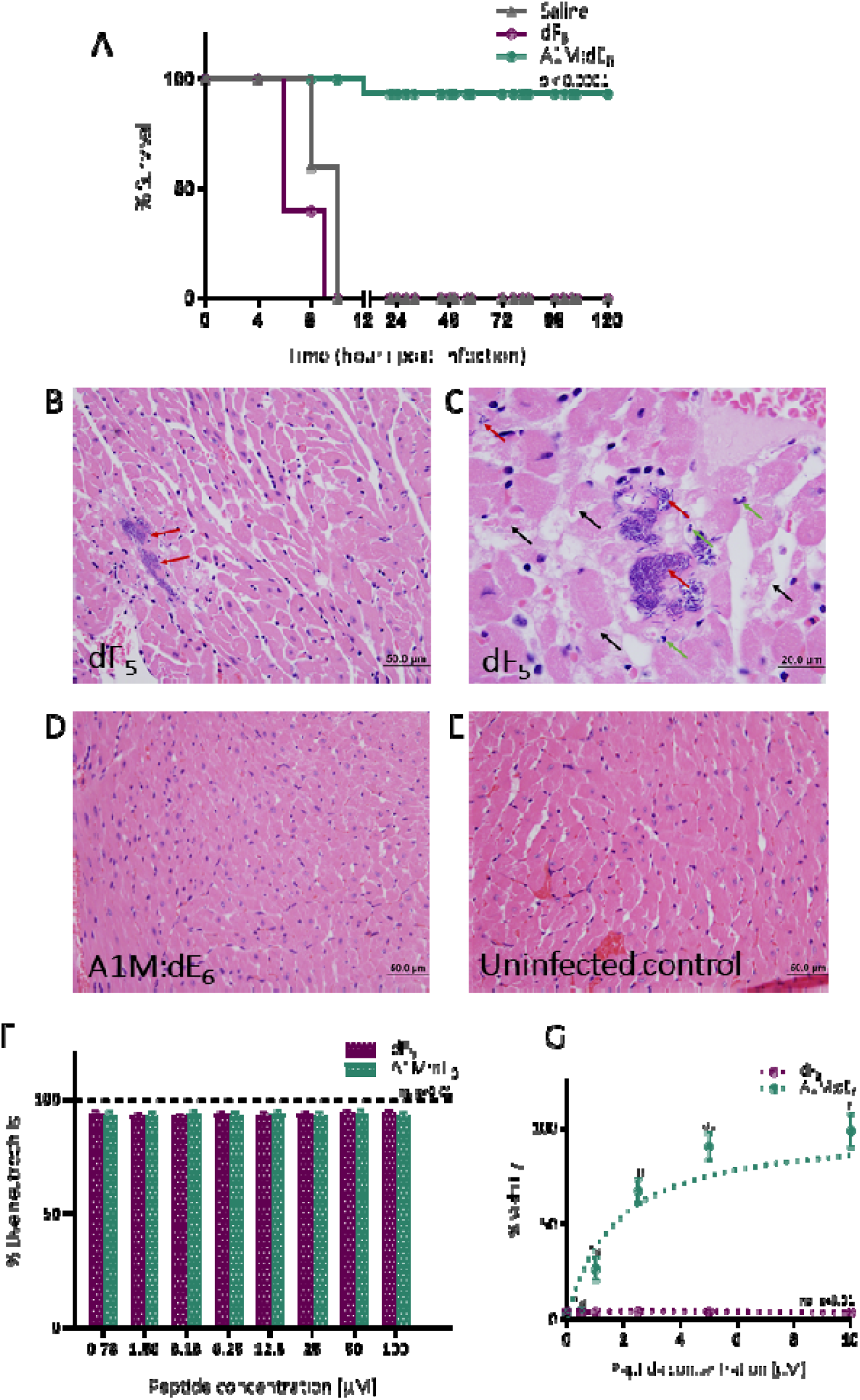
PapR_7_ – A1M:dE_6_ enhances protective immunity against bacteremia infection. (A) Survival of mice in a bacteremia infection caused by the Bc ATCC 14579 strain (n=15). p < 0.0001 when comparing the mortality of infected mice treated with PapR_7_ – A1M:dE_6_ against those treated with sterile saline by using the Kaplan−Meier Log Rank (Mantel−Cox) survival test. (B-D) Representative histopathological images of mouse heart infected by Bc treated with (B-C) PapR_7_ –dF_5_ or (D) PapR_7_ – A1M:dE_6_. (E) Uninfected control (H&E staining; B, D & E: 400x, C :1000x; Black arrows, myocardial necrosis and cardiomyolysis. Red arrows, presence of bacterial microcolonies. Green arrows, scattered neutrophils). (F) Cytotoxicity of PapR_7_ – A1M:dE_6_ and dF_5_ on mouse neutrophils (mean ± SEM, n = 9). ns indicates no statistical difference between treatments and different concentration of examined peptides. (G) Neutrophil viability after challenge with supernatants from Bc cultures treated with PapR_7_ – A1M:dE_6_ or dF_5_ (0.5-10μM, mean ± SEM, n = 9). *p < 0.01 indicates a statistically significant difference between untreated Bc cultures supernatants versus those treated with PapR_7_ – A1M:dE_6_. Different letters indicate statistically significant differences between supernatants from Bc cultures treated with different concentrations of PapR_7_ – A1M:dE_6_ (p < 0.01). ns indicates no statistical difference.

Neutrophils are an essential component of the innate immune system, as they are the first leukocytes to penetrate and accumulate rapidly in the infected tissues (Amulic et al., 2012; Livingston et al., 2019). To better understand our *in-vivo* results and to further support out hypothesis that neutrophils in addition to macrophages are being eliminated by the QS secreted virulence factors, we isolated neutrophils from mouse bone marrow and verified whether the protective effect of PapR7 – A1M:dE6 remained the same as observed for macrophages. After determining that none of the examined peptides was cytotoxic to polymorphonuclear neutrophils (PMNs) cells (Figure 3E), we assessed their viability in response to Bc culture supernatants treated with either PapR_7_ – A1M:dE_6_ or PapR_7_ –dF_5_ (Figure 3F). PMNs were highly susceptible to Bc supernatants grown with or without PapR_7_ –dF_5_, with almost 100% loss in cell viability. However, when challenged with supernatants treated with PapR_7_ – A1M:dE_6_, their viability was protected in a concentration dependent manner (Figure 3F). These results concur with our hypothesis on the possible role of PapR_7_ – A1M:dE_6_ in preserving host innate immune responses mediated by macrophages and neutrophils against Bc infection. Given that PapR_7_ – A1M:dE_6_ exposure inhibits the production of PlcR-dependent virulence factors *in-vitro* (Figure 1B, D) and the high survival rates *in-vivo* suggests that PapR_7_ – A1M:dE_6_ may promotes host immune cells rescue. To the best of our knowledge this is the first evidence that demonstrate the contribution of QS system in impairing the innate immune system activity that allows the pathogen to proliferate rapidly.

### PapR_7_ – A1M:dE_6_ blocks Bc host immune system eradication

Finally, to better understand the importance of host neutrophils protection by PapR_7_ – A1M:dE_6_ inhibitory activity, we examined Bc infection in neutropenic mouse lung and thigh, as previously described (Cheah et al., 2015; Garcia Chavez et al., 2021). Mice were rendered neutropenic by the administration of cyclophosphamide, and then both acute pneumonia and thigh infections were performed as we have previously shown (Figure 1E, S1D). When these mice were inoculated with Bc, they were too susceptible to lung infection and died within 6 hours after challenge, suggesting that depletion of neutrophils make the mice more vulnerable to lung infection and respiratory arrest by Bc. Therefore, we decided to euthanize mice from all groups for bacterial burden determination after only one treatment (2 hpi) with the tested peptides (PapR_7_ – A1M:dE_6_ or dF_5_). It is interesting to see that despite its significant reduction in bacterial counts during acute pneumonia (Figure 1E), PapR_7_ – A1M:dE_6_ failed to attenuated Bc burden in neutropenic mouse lungs (Figure 4A). To further support this observation, we performed the neutropenic mouse thigh infection model and still, no significant different was observed in thigh burden of Bc with or without PapR_7_ – A1M:dE_6_ treatment after 24 h (p < 0.01; Figure S4A). Collectively, these results indicate that PapR_7_ – A1M:dE_6_ has a role in protecting the host immune cells, and when these cells were depleted by Bc, the protective effect of PapR_7_ – A1M:dE_6_ peptide is lost. Because PapR_7_ – A1M:dE_6_ peptide is an non-bactericidal anti–virulent agent, we hypothesized that the repression of PlcR-dependent virulence factors by PapR_7_ – A1M:dE_6_ peptide prevents the killing of the host immune cells (macrophages and neutrophils), and thus enable them to promote clearance of Bc. In order to identify the toxic PlcR-dependent components, we evaluated the different composition in the secretome (i.e protein released in the culture medium) of Bc that was grown with and without PapR_7_ – A1M:dE_6_. The bacterial supernatants were filtered, precipitated with TCA, and analyzed using SDS-PAGE (Figure 4B). Both native and PapR_7_ –dF_5_ treatments showed similarities in their secretome profiles, and contained significant amounts of secreted proteins (Figure 4B, first and second lane). In contrast, the secretome profile of Bc culture treated with PapR_7_ – A1M:dE_6_ showed significantly reduction of more than five proteins (Figure 4B, third lane). To further characterize these secreted virulence proteins, we carried out a proteomic analysis of the bacterial secretomes using LC-MS/MS analysis and label-free quantification (LFQ; Figure 4C). We determined which virulence factors are differently secreted by calculating the normalized protein abundances of all secreted proteins in each treatment based on their LFQ intensity values. Difference of at least 2-fold in LFQ with p < 0.05 values were considered significant. Exposure of PapR_7_ – A1M:dE_6_ led to a significant decrease in abundances of three common proteins associated with membrane-damaging and cell lysis; the binding component of Hemolysin BL (Hbl-B), and the two phosphatidylcholine esterases-Sphingomyelinase (SMase) and Phospholipase C, which together form cereolysin AB (Figure 4A; Gilmore et al., 1989; Visiello et al., 2016). Secretion of these toxins was 4.5-6 log-fold reduced in Bc treated with PapR_7_ – A1M:dE_6_. However, no significant different in levels of secretion was observed between native or PapR_7_–dF_5_ treated bacterial secretomes.

**Figure 4:**
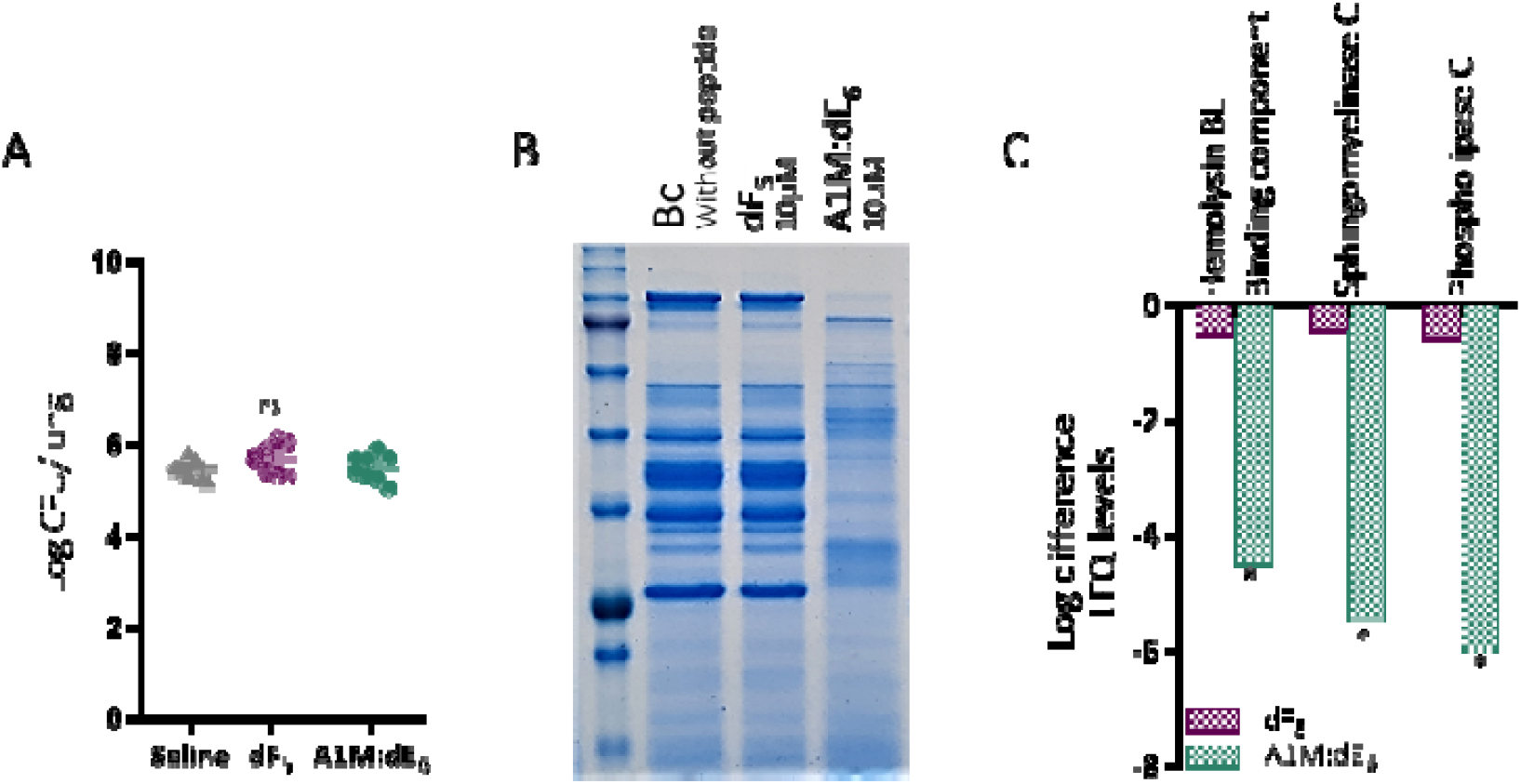
PapR_7_ – A1M:dE6 block Bc immune escape. (A) Bacterial loads in the lung from neutropenic mice post-infection with saline, PapR_7_–dF_5_ and A1M:dE_6_ (mean ± SEM, n = 8). ns indicates no statistically significant difference between treatments. (B) SDS-PAGE analysis showing secretome profiles from Bc strain treated with 10μM PapR_7_ – A1M:dE_6_ or dF_5._ The gel is representative for three biologically independent experiments. (C) Log difference in normalized LFQ values determined for proteins released of Bc strain treated with 10μM PapR_7_ – A1M:dE_6_ or dF_5_. Statistical analysis (n=3) was performed using the Perseus statistical package. *p < 0.05 indicates a statistically significant difference between untreated Bc and addition of PapR_7_ – A1M:dE_6_.

Previous studies have shown that these exotoxins contributed to the pathogenicity of Bc during infections through positively regulation of PlcR (Enosi Tuipulotu et al., 2021; Salamitou et al., 2000; Sastalla et al., 2013). These studies along with our results shed light on a novel mechanism that enable PapR_7_ – A1M:dE6 to attenuate Bc infections. Its ability to significantly reduce the secretion of at least three PlcR-regulated major toxins protects the host immune cells from eradication and thusenhance host defense.

## Discussion

Evading host innate defenses play a key role in pathogenicity of opportunistic bacteria, and determines the outcome of infection. Professional phagocytes, such as macrophages, neutrophils, dendritic and mast cells constitute the first lines of host defense against infectious agents (Lehman and Segal, 2020; Netea et al., 2011). Some pathogens evade from these phagocytes recognition by modulating their pathogen associated molecular patterns (PAMPs), hiding and replicating intracellularly, or killing them directly through secreted toxin and/or indirectly by inducing apoptosis (Kaufmann and Dorhoi, 2016; Leseigneur et al., 2020; Tran and Ramarao, 2013). The discovery that a wide spectrum of pathogens utilize QS to regulate virulence traits makes it an attractive target for the development of alternative anti-infectives. If virulence can be controlled, then it is suggested that the host immune system will be capable of overcoming any invading bacteria. The PapR-PlcR QS system of Bc regulates the expression of virulence determinants during host infection (Beecher et al., 2000; Enosi Tuipulotu et al., 2021; Salamitou et al., 2000), therefore, could equip Bc with immune evading machinery when sufficient bacterial threshold is amassed. By doing so, a concerted attack is achieved, to overwhelm host defense mechanisms and establish infection. However, the role of the regulator PlcR in establishing Bc infection or the molecular strategies allowing the pathogen to resist host immune defense system have been elusive. By using QS null mutant strains, several studies have shown that Bc escapes the immune system independent of the PlcR regulator. For examples, Bc spores survive, germinate and escape from macrophages (Ramarao and Lereclus, 2005) eventually leading to the production of secreted toxin HlyII responsible for host cell death (Tran et al., 2011). Despite remarkable insight into the molecular mechanisms driving Bc pathogenesis, we cannot exclude the possibility for other unknown factors.

Previously, we reported on the first PapR_7_-derived peptidic inhibitors of Bc PlcR-PapR QS system with high potential to serve as a novel systemically administered QQ agents (Yehuda et al., 2018, 2019). However, their potential utility in attenuating virulence during host infection remain unclear. In this study, we reveal the most potent peptidic inhibitor of Bc QS system to be reported so far, PapR_7_ – A1M:dE_6_, has the ability to suppress the expression of PlcR-mediated virulence factors, as reflected by a loss in hemolytic activity. Furthermore, PapR_7_ – A1M:dE_6_ reduces bacterial burden in both thigh and acute lung infections in mice, demonstrating its role in promoting bacterial clearance.

Prior studies have explored QQ compounds for their therapeutic potential against Gram-positive bacterial infections. Most of these studies have only demonstrated antibacterial effect but failed to provide clear molecular mechanisms exclusively to QQ underlying their therapeutic effect. To prove that PapR_7_ – A1M:dE_6_ intercepts the interplay between bacterial virulence factors and host immune responses; we focused on the interactions between Bc and innate immune cells. We demonstrated that PapR_7_ – A1M:dE_6_ confers protective effect to macrophages and neutrophils *in-vitro*, by rescuing them from BC-mediated cytoxicity by inhibiting the expression of PlcR-regulated Hbl-B and cereolysin AB. More significantly, PapR_7_ – A1M:dE_6_ was efficacious in attenuating mortality in a mouse model of bacteremia. The importance of neutrophils is further demonstrated by the observation that PapR_7_ – A1M:dE_6_ failed to attenuate Bc burden in lung and thigh of immunosuppressed neutropenic mice. To the best of our knowledge, this is the first reported demonstration of the QS system in impairing the innate immune system during bacterial infection, and thus emphasizes the therapeutic potential of PapR_7_ – A1M:dE_6_ and other QQ agents through their protective roles for components of immune systems such as macrophages and neutrophils. Using our synthetic PapR_7_ – A1M:dE_6_ inhibitor and comprehensive analysis, we uncovered at least three of PlcR-regulated major toxins underlying Bc pathogenicity during host infection, such as Hemolysin BL (Hbl-B), and the two phosphatidylcholine esterases-Sphingomyelinase (SMase) and Phospholipase C, which together form cereolysin AB.

Taken together, our study reveal that the PapR_7_ – A1M:dE_6_ can act as a novel immune-therapeutic agent to combat Bc infections. Successful implementation either as the sole agent or in combination with current antimicrobial agents or antibiotics could attenuate Bc virulence without inducing strong selective pressure for resistance development. Most importantly, our synthetic PapR_7_-derived inhibitor demonstrates for the first time, that PlcR-PapR QS-regulated Bc pathogenesis significantly modulates host immunity. By significantly reducing the secretion of PlcR-regulated virulence factors, PapR_7_ – A1M:dE_6_ protects the host immune cells from eradication and thus enhance host defense. These peptidic QQ agents and the insights that they provide constitute new and readily accessible chemical tools for studying other QS systems and how they modulate the host response. Moreover, they could lead to the development of new chemical strategies to attenuate variety of infectious diseases.

## Notes

### Competing Interest Statement

The authors have declared no competing interest.

## References

Agaisse, H., Gominet, M., Okstad, O. a, Kolstø, a B., and Lereclus, D. (1999). PlcR is a pleiotropic regulator of extracellular virulence factor gene expression in Bacillus thuringiensis. Molecular Microbiology 32, 1043–1053. https://doi.org/10.1046/j.1365-2958.1999.01419.x.

Allen, R.C., Popat, R., Diggle, S.P., and Brown, S.P. (2014). Targeting virulence: Can we make evolution-proof drugs? Nature Reviews Microbiology 12, 300–308. https://doi.org/10.1038/nrmicro3232.

Amulic, B., Cazalet, C., Hayes, G.L., Metzler, K.D., and Zychlinsky, A. (2012). Neutrophil Function: From Mechanisms to Disease. Annual Review of Immunology 30, 459–489. https://doi.org/10.1146/annurev-immunol-020711-074942.

Bassler, B.L., and Losick, R. (2006). Bacterially Speaking. Cell 125, 237–246. https://doi.org/10.1016/j.cell.2006.04.001.

Beecher, D.J., and Wong, A.C.L. (2000). Cooperative, synergistic and antagonistic haemolytic interactions between haemolysin BL, phosphatidylcholine phospholipase C and sphingomyelinase from Bacillus cereus. Microbiology 146, 3033–3039. https://doi.org/10.1099/00221287-146-12-3033.

Beecher, D.J., Olsen, T.W., Somers, E.B., and Wong, A.C.L. (2000). Evidence for contribution of tripartite hemolysin BL, phosphatidylcholine-preferring phospholipase C, and collagenase to virulence of Bacillus cereus endophthalmitis. Infection and Immunity 68, 5269–5276. https://doi.org/10.1128/IAI.68.9.5269-5276.2000.

Bottone, E.J. (2010). Bacillus cereus, a volatile human pathogen. Clinical Microbiology Reviews 23, 382–398. https://doi.org/10.1128/CMR.00073-09.

Bouillaut, L., Perchat, S., Arold, S., Zorrilla, S., Slamti, L., Henry, C., Gohar, M., Declerck, N., and Lereclus, D. (2008). Molecular basis for group-specific activation of the virulence regulator PlcR by PapR heptapeptides. Nucleic Acids Research 36, 3791–3801. https://doi.org/10.1093/nar/gkn149.

C., W. (2000). Molecular mechanisms that confer antibacterial drug resistance. Nature 406, 775–781..

Case, R.J., Labbate, M., and Kjelleberg, S. (2008). AHL-driven quorum-sensing circuits: Their frequency and function among the Proteobacteria. ISME Journal 2, 345–349. https://doi.org/10.1038/ismej.2008.13.

Cheah, S.E., Wang, J., Nguyen, V.T. h. T., Turnidge, J.D., Li, J., and Nation, R.L. (2015). New pharmacokinetic/pharmacodynamic studies of systemically administered colistin against Pseudomonas aeruginosa and Acinetobacter baumannii in mouse thigh and lung infection models: smaller response in lung infection. The Journal of Antimicrobial Chemotherapy 70, 3291–3297. https://doi.org/10.1093/jac/dkv267.

Cox, J., Hein, M.Y., Luber, C.A., Paron, I., Nagaraj, N., and Mann, M. (2014). Accurate proteome-wide label-free quantification by delayed normalization and maximal peptide ratio extraction, termed MaxLFQ. Molecular and Cellular Proteomics 13, 2513–2526. https://doi.org/10.1074/mcp.M113.031591.

Dunny, G.M., and Leonard, B.A. (1997). Cell-cell communication in gram-positive bacteria. Annual Review of Microbiology 51, 527–564. https://doi.org/10.1146/annurev.micro.51.1.527.

Enosi Tuipulotu, D., Mathur, A., Ngo, C., and Man, S.M. (2021). Bacillus cereus: Epidemiology, Virulence Factors, and Host–Pathogen Interactions. Trends in Microbiology 29, 458–471. https://doi.org/10.1016/j.tim.2020.09.003.

Galloway, W.R.J.D., Hodgkinson, J.T., Bowden, S., Welch, M., and Spring, D.R. (2012). Applications of small molecule activators and inhibitors of quorum sensing in Gram-negative bacteria. Trends in Microbiology 20, 449–458. https://doi.org/10.1016/j.tim.2012.06.003.

Garcia Chavez, M., Garcia, A., Lee, H.Y., Lau, G.W., Parker, E.N., Komnick, K.E., and Hergenrother, P.J. (2021). Synthesis of Fusidic Acid Derivatives Yields a Potent Antibiotic with an Improved Resistance Profile. ACS Infectious Diseases 7, 493–505. https://doi.org/10.1021/acsinfecdis.0c00869.

Gilmore, M.S., Cruz-Rodz, A.L., Leimeister-Wachter, M., Kreft, J., and Goebel, W. (1989). A Bacillus cereus cytolytic determinant, cereolysin AB, which comprises the phospholipase C and sphingomyelinase genes: Nucleotide sequence and genetic linkage. Journal of Bacteriology 171, 744–753. https://doi.org/10.1128/jb.171.2.744-753.1989.

Gilois, N., Ramarao, N., Bouillaut, L., Perchat, S., Aymerich, S., Nielsen-LeRoux, C., Lereclus, D., and Gohar, M. (2007). Growth-related variations in the Bacillus cereus secretome. Proteomics 7, 1719–1728. https://doi.org/10.1002/pmic.200600502.

Glasset, B., Herbin, S., Granier, S.A., Cavalie, L., Lafeuille, E., Guérin, C., Ruimy, R., Casagrande-Magne, F., Levast, M., Chautemps, N., et al. (2018). Bacillus cereus, a serious cause of nosocomial infections: Epidemiologic and genetic survey. PLoS ONE 13, 1–19. https://doi.org/10.1371/journal.pone.0194346.

Gohar, M., Økstad, O.A., Gilois, N., Sanchis, V., Kolstø, A.B., and Lereclus, D. (2002). Two-dimensional electrophoresis analysis of the extracellular proteome of Bacillus cereus reveals the importance of the PlcR regulon. In Proteomics, p.

Gohar, M., Faegri, K., Perchat, S., Ravnum, S., Økstad, O.A., Gominet, M., Kolstø, A.B., and Lereclus, D. (2008). The PlcR virulence regulon of Bacillus cereus. PLoS ONE 3. https://doi.org/10.1371/journal.pone.0002793.

Gominet, M., Slamti, L., Gilois, N., Rose, M., and Lereclus, D. (2001). Oligopeptide permease is required for expression of the Bacillus thuringiensis plcR regulon and for virulence. Molecular Microbiology 40, 963–975. https://doi.org/10.1046/j.1365-2958.2001.02440.x.

Grandclément, C., Tannières, M., Moréra, S., Dessaux, Y., and Faure, D. (2015). Quorum quenching: Role in nature and applied developments. FEMS Microbiology Reviews https://doi.org/10.1093/femsre/fuv038.

Granum, P.E., and Lund, T. (1997). Bacillus cereus and its food poisoning toxins. FEMS Microbiology Letters 157, 223–228..

Grenha, R., Slamti, L., Nicaise, M., Refes, Y., Lereclus, D., and Nessler, S. (2013). Structural basis for the activation mechanism of the PlcR virulence regulator by the quorum-sensing signal peptide PapR. Proceedings of the National Academy of Sciences 110, 1047–1052. https://doi.org/10.1073/pnas.1213770110.

Helgason, E., Okstad, O.A., Caugant, D.A., Johansen, H.A., Fouet, A., Mock, M., Hegna, I., and Kolstø, A.B. (2000). Bacillus anthracis, Bacillus cereus, and Bacillus thuringiensis--one species on the basis of genetic evidence. Applied and Environmental Microbiology 66, 2627–2630. https://doi.org/10.1128/AEM.66.6.2627-2630.2000.

Hernandez, E., Ramisse, F., Ducoureau, J.P., Cruel, T., and Cavallo, J.D. (1998). Bacillus thuringiensis subsp. konkukian (Serotype H34) superinfection: Case report and experimental evidence of pathogenicity in immunosuppressed mice. Journal of Clinical Microbiology 36, 2138–2139. https://doi.org/10.1128/jcm.36.7.2138-2139.1998.

Ivanova, N., Sorokin, A., Anderson, I., Galleron, N., Candelon, B., Kapatral, V., Bhattacharyya, A., Reznik, G., Mikhailova, N., Lapidus, A., et al. (2003). Genome sequence of Bacillus cereus and comparative analysis with Bacillus anthracis. Nature https://doi.org/10.1038/nature01582.

Kaufmann, S.H.E., and Dorhoi, A. (2016). Molecular Determinants in Phagocyte-Bacteria Interactions. Immunity 44, 476–491. https://doi.org/10.1016/j.immuni.2016.02.014.

Lehman, H.K., and Segal, B.H. (2020). The role of neutrophils in host defense and disease. Journal of Allergy and Clinical Immunology 145, 1535–1544. https://doi.org/10.1016/j.jaci.2020.02.038.

Lereclus, D., Arantès, O., Chaufaux, J., and Lecadet, M. (1989). Transformation and expression of a cloned delta-endotoxin gene in Bacillus thuringiensis. FEMS Microbiol Lett 51, 211–217..

Lereclus, D., Agaisse, H., Gominet, M., Salamitou, S., and Sanchis, V. (1996). Identification of a Bacillus thuringiensis gene that positively regulates transcription of the phosphatidylinositol-specific phospholipase C gene at the onset of the stationary phase. Journal of Bacteriology 178, 2749–2756. https://doi.org/10.1128/jb.178.10.2749-2756.1996.

Leseigneur, C., Lê-Bury, P., Pizarro-Cerdá, J., and Dussurget, O. (2020). Emerging Evasion Mechanisms of Macrophage Defenses by Pathogenic Bacteria. Frontiers in Cellular and Infection Microbiology 10, 1–9. https://doi.org/10.3389/fcimb.2020.577559.

Liu, J., Zuo, Z., Sastalla, I., Liu, C., Jang, J.Y., Sekine, Y., Li, Y., Pirooznia, M., Leppla, S.H., Finkel, T., et al. (2020). Sequential CRISPR-Based Screens Identify LITAF and CDIP1 as the Bacillus cereus Hemolysin BL Toxin Host Receptors. Cell Host and Microbe 28, 402-410.e5. https://doi.org/10.1016/j.chom.2020.05.012.

Livingston, E.T., Mursalin, M.H., and Callegan, M.C. (2019). A pyrrhic victory: The PMN response to ocular bacterial infections. Microorganisms 7. https://doi.org/10.3390/microorganisms7110537.

Mattmann, M.E., and Blackwell, H.E. (2010). Small molecules that modulate quorum sensing and control virulence in pseudomonas aeruginosa. Journal of Organic Chemistry 75, 6737–6746. https://doi.org/10.1021/jo101237e.

McBrayer, D.N., Gantman, B.K., Cameron, C.D., and Tal-Gan, Y. (2017). An Entirely Solid Phase Peptide Synthesis-Based Strategy for Synthesis of Gelatinase Biosynthesis-Activating Pheromone (GBAP) Analogue Libraries: Investigating the Structure–Activity Relationships of the Enterococcus faecalis Quorum Sensing Signal. Organic Letters 19, 3295–3298. https://doi.org/10.1021/acs.orglett.7b01444.

McBrayer, D.N., Cameron, C.D., Gantman, B.K., and Tal-Gan, Y. (2018). Rational Design of Potent Activators and Inhibitors of the Enterococcus faecalis Fsr Quorum Sensing Circuit. ACS Chemical Biology 13, 2673–2681. https://doi.org/10.1021/acschembio.8b00610.

McBrayer, D.N., Cameron, C.D., and Tal-Gan, Y. (2020). Development and utilization of peptide-based quorum sensing modulators in Gram-positive bacteria. Organic and Biomolecular Chemistry 18, 7273–7290. https://doi.org/10.1039/d0ob01421d.

McGill, C.J., Lu, R.J., and Benayoun, B.A. (2021). Protocol for analysis of mouse neutrophil NETosis by flow cytometry. STAR Protocols 2, 100948. https://doi.org/10.1016/j.xpro.2021.100948.

Miller, M.B., and Bassler, B.L. (2001). Quorum Sensing in Bacteria. Annual Review of Microbiology 55, 165–199. https://doi.org/10.1146/annurev.micro.55.1.165.

Nakayama, J., Yokohata, R., Sato, M., Suzuki, T., Matsufuji, T., Nishiguchi, K., Kawai, T., Yamanaka, Y., Nagata, K., Tanokura, M., et al. (2013). Development of a Peptide Antagonist against fsr Quorum Sensing of Enterococcus faecalis. ACS Chemical Biology 8, 804–811. https://doi.org/10.1021/cb300717f.

Neiditch, M.B., Capodagli, G.C., Prehna, G., and Federle, M.J. (2017). Genetic and Structural Analyses of RRNPP Intercellular Peptide Signaling of Gram-Positive Bacteria. Annual Review of Genetics 51, 311–333. https://doi.org/10.1146/annurev-genet-120116-023507.

Netea, M.G., Quintin, J., and Van Der Meer, J.W.M. (2011). Trained immunity: A memory for innate host defense. Cell Host and Microbe 9, 355–361. https://doi.org/10.1016/j.chom.2011.04.006.

Papenfort, K., and Bassler, B.L. (2016). Quorum sensing signal-response systems in Gram-negative bacteria. Nature Reviews Microbiology 14, 576–588. https://doi.org/10.1038/nrmicro.2016.89.

Piewngam, P., Chiou, J., Chatterjee, P., and Otto, M. (2020). Alternative approaches to treat bacterial infections: targeting quorum-sensing. Expert Review of Anti-Infective Therapy 18, 499–510. https://doi.org/10.1080/14787210.2020.1750951.

Pomerantsev, A.P., Kalnin, K. V., Osorio, M., and Leppla, S.H. (2003). Phosphatidylcholine-Specific Phospholipase C and Sphingomyelinase Activities in Bacteria of the Bacillus cereus Group. Infection and Immunity 71, 6591–6606. https://doi.org/10.1128/IAI.71.11.6591-6606.2003.

Rajput, A., Kaur, K., and Kumar, M. (2016). SigMol: Repertoire of quorum sensing signaling molecules in prokaryotes. Nucleic Acids Research https://doi.org/10.1093/nar/gkv1076.

Ramarao, N., and Lereclus, D. (2005). The InhA 1 metalloprotease allows spores of the B. cereus group to escape macrophages. Cellular Microbiology 7, 1357–1364. https://doi.org/10.1111/j.1462-5822.2005.00562.x.

Ramarao, N., and Sanchis, V. (2013). The pore-forming haemolysins of Bacillus cereus: A review. Toxins https://doi.org/10.3390/toxins5061119.

Rasko, D.A., and Sperandio, V. (2010). Anti-virulence strategies to combat bacteria-mediated disease. Nature Reviews Drug Discovery 9, 117–128. https://doi.org/10.1038/nrd3013.

Rasko, D.A., Altherr, M.R., Han, C.S., and Ravel, J. (2005). Genomics of the Bacillus cereus group of organisms. FEMS Microbiology Reviews https://doi.org/10.1016/j.femsre.2004.12.005.

Rutherford, S.T., and Bassler, B.L. (2012). Bacterial quorum sensing: its role in virulence and possibilities for its control. Cold Spring Harbor Perspectives in Medicine 2. https://doi.org/10.1101/cshperspect.a012427.

Salamitou, S., Ramisse, F., Brehelin, M., Bourguet, D., Gilois, N., Gominet, M., Hernandez, E., and Lereclus, D. (2000). The plcR regulon is involved in the opportunistic properties of Bacillus thuringiensis and Bacillus cereus in mice and insects. Microbiology 146, 2825–2832. https://doi.org/10.1099/00221287-146-11-2825.

Sastalla, I., Fattah, R., Coppage, N., Nandy, P., Crown, D., Pomerantsev, A.P., and Leppla, S.H. (2013). The Bacillus cereus Hbl and Nhe Tripartite Enterotoxin Components Assemble Sequentially on the Surface of Target Cells and Are Not Interchangeable. PLoS ONE 8. https://doi.org/10.1371/journal.pone.0076955.

Scheltema, R.A., Hauschild, J.P., Lange, O., Hornburg, D., Denisov, E., Damoc, E., Kuehn, A., Makarov, A., and Mann, M. (2014). The Q exactive HF, a benchtop mass spectrometer with a pre-filter, high-performance quadrupole and an ultra-high-field orbitrap analyzer. Molecular and Cellular Proteomics 13, 3698–3708. https://doi.org/10.1074/mcp.M114.043489.

Slamti, L., and Lereclus, D. (2002). A cell-cell signaling peptide activates the PlcR virulence regulon in bacteria of the Bacillus cereus group. EMBO Journal 21, 4550–4559. https://doi.org/10.1093/emboj/cdf450.

Slamti, L., Perchat, S., Gominet, M., Vilas-Bôas, G., Fouet, A., Mock, M., Sanchis, V., Chaufaux, J., Gohar, M., and Lereclus, D. (2004). Distinct mutations in PlcR explain why some strains of the Bacillus cereus group are nonhemolytic. Journal of Bacteriology https://doi.org/10.1128/JB.186.11.3531-3538.2004.

Slamti, L., Perchat, S., Huillet, E., and Lereclus, D. (2014a). Quorum sensing in Bacillus thuringiensis is required for completion of a full infectious cycle in the insect. Toxins 6, 2239–2255. https://doi.org/10.3390/toxins6082239.

Slamti, L., Perchat, S., Huillet, E., and Lereclus, D. (2014b). Quorum sensing in Bacillus thuringiensis is required for completion of a full infectious cycle in the insect. Toxins 6, 2239–2255. https://doi.org/10.3390/toxins6082239.

Stenfors Arnesen, L.P., Fagerlund, A., and Granum, P.E. (2008). From soil to gut: Bacillus cereus and its food poisoning toxins. FEMS Microbiology Reviews 32, 579–606. https://doi.org/10.1111/j.1574-6976.2008.00112.x.

Swamydas, M., and Lionakis, M.S. (2013). Isolation, Purification and Labeling of Mouse Bone Marrow Neutrophils for Functional Studies and Adoptive Transfer Experiments. Journal of Visualized Experiments 1–7. https://doi.org/10.3791/50586.

Tal-Gan, Y., Ivancic, M., Cornilescu, G., Cornilescu, C.C., and Blackwell, H.E. (2013a). Structural characterization of native autoinducing peptides and abiotic analogues reveals key features essential for activation and inhibition of an agrc quorum sensing receptor in staphylococcus aureus. Journal of the American Chemical Society 135, 18436–18444. https://doi.org/10.1021/ja407533e.

Tal-Gan, Y., Stacy, D.M., Foegen, M.K., Koenig, D.W., and Blackwell, H.E. (2013b). Highly potent inhibitors of quorum sensing in staphylococcus aureus revealed through a systematic synthetic study of the group-III autoinducing peptide. Journal of the American Chemical Society 135, 7869–7882. https://doi.org/10.1021/ja3112115.

Tran, S.L., and Ramarao, N. (2013). Bacillus cereus immune escape: A journey within macrophages. FEMS Microbiology Letters https://doi.org/10.1111/1574-6968.12209.

Tran, S.L., Guillemet, E., Gohar, M., Lereclus, D., and Ramarao, N. (2010). CwpFM (EntFM) is a Bacillus cereus potential cell wall peptidase implicated in adhesion, biofilm formation, and virulence. Journal of Bacteriology 192, 2638–2642. https://doi.org/10.1128/JB.01315-09.

Tran, S.L., Guillemet, E., Ngo-Camus, M., Clybouw, C., Puhar, A., Moris, A., Gohar, M., Lereclus, D., and Ramarao, N. (2011). Haemolysin II is a Bacillus cereus virulence factor that induces apoptosis of macrophages. Cellular Microbiology 13, 92–108. https://doi.org/10.1111/j.1462-5822.2010.01522.x.

Uroz, S., Dessaux, Y., and Oger, P. (2009). Quorum sensing and quorum quenching: The Yin and Yang of bacterial communication. ChemBioChem 10, 205–216. https://doi.org/10.1002/cbic.200800521.

Utari, P.D., Setroikromo, R., Melgert, B.N., and Quax, W.J. (2018). PvdQ quorum quenching acylase attenuates Pseudomonas aeruginosa virulence in a mouse model of pulmonary infection. Frontiers in Cellular and Infection Microbiology 8. https://doi.org/10.3389/fcimb.2018.00119.

Visiello, R., Colombo, S., and Carretto, E. (2016). Chapter 3 - Bacillus cereus Hemolysins and Other Virulence Factors. V.B.T.-T.D.F. of B. tcereus Savini, ed. (Academic Press), pp. 35–44.

Welsh, M.A., and Blackwell, H.E. (2016). Chemical probes of quorum sensing: From compound development to biological discoverya. FEMS Microbiology Reviews 40, 774–794. https://doi.org/10.1093/femsre/fuw009.

Yang, Y., Koirala, B., Sanchez, L.A., Phillips, N.R., Hamry, S.R., and Tal-Gan, Y. (2017). Structure-Activity Relationships of the Competence Stimulating Peptides (CSPs) in Streptococcus pneumoniae Reveal Motifs Critical for Intra-group and Cross-group ComD Receptor Activation. ACS Chemical Biology https://doi.org/10.1021/acschembio.7b00007.

Yang, Y., Lin, J., Harrington, A., Cornilescu, G., Lau, G.W., and Tal-Gan, Y. (2020). Designing cyclic competence-stimulating peptide (CSP) analogs with pan-group quorum-sensing inhibition activity in Streptococcus pneumoniae. Proceedings of the National Academy of Sciences of the United States of America 117, 1689–1699. https://doi.org/10.1073/pnas.1915812117.

Yehuda, A., Slamti, L., Bochnik-Tamir, R., Malach, E., Lereclus, D., and Hayouka, Z. (2018). Turning off Bacillus cereus quorum sensing system with peptidic analogs. Chemical Communications https://doi.org/10.1039/c8cc05496g.

Yehuda, A., Slamti, L., Malach, E., Lereclus, D., and Hayouka, Z. (2019). Elucidating the Hot Spot Residues of Quorum Sensing Peptidic Autoinducer PapR by Multiple Amino Acid Replacements. Frontiers in Microbiology https://doi.org/10.3389/fmicb.2019.01246.

Zhu, L., and Lau, G.W. (2011). Inhibition of competence development, horizontal gene transfer and virulence in streptococcus pneumoniae by a modified competence stimulating peptide. PLoS Pathogens 7. https://doi.org/10.1371/journal.ppat.1002241.

