## Supporting information for "Quorum sensing peptidic inhibitor rescue host immune system eradication: a novel QS infectivity mechanism"

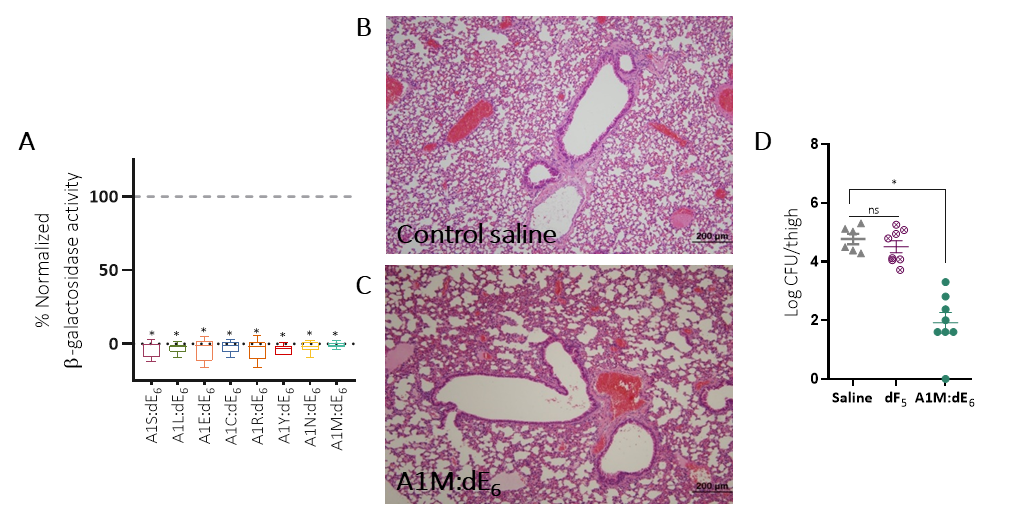
***Supplementary data***

**Figure S1:** **PapR_7_ – A1M:dE_6_ attenuates Bc pathogenesis during acute lung infection, related to Figure 1.** (A) β-galactosidase activity of Bt ΔpapR A'Z induced by the addition of 10 µM A1-substituted:dE_6_ peptides normalized to synthetic PapR_7_ peptide at onset of the stationary phase of bacterial growth (OD_600_ of 3 ± 0.5, mean ± SEM, n = 9). ^*^p ˂ 0.01 when comparing between synthetic PapR_7_ peptide and A1-substituted:dE_6_ peptides. (B-C) Histopathological analysis of lungs in saline control group versus PapR_7_ – A1M:dE_6_ treated mice (100x). (D) Thigh bacterial load from mice post Bc infection treated with saline, PapR_7_–dF_5_ and A1M:dE_6_ (mean ± SEM, n = 6-8). ^∗^p < 0.01 indicates a statistically significant difference between treatments. ns indicates no statistically significant difference between treatments.

**Table S1:** Comparison between IC_50_ values of new A1-substituted:dE_6_ peptides and their parent inhibitor PapR_7_ –dE_6_, as determined by the lacZ-based reporter assay.

| IC_50_ [µM] | PlcR activation | Sequence | | | | | | | Peptide name |
| --- | --- | --- | --- | --- | --- | --- | --- | --- | --- |
| 0.977 ± 0.04 | - | F | dE | F | P | L | D | A | **PapR_7_ - dE_6_** |
| 2.569 ± 0.343 | **-** | F | dE | F | P | L | D | S | **PapR_7_ - A1S:dE_6_** |
| 0.752 ± 0.111 | **-** | F | dE | F | P | L | D | L | **PapR_7_ - A1L:dE_6_** |
| **-** | **-** | F | dE | F | P | L | D | E | **PapR_7_ - A1E:dE_6_** |
| **-** | **-** | F | dE | F | P | L | D | C | **PapR_7_ - A1C:dE_6_** |
| **-** | **-** | F | dE | F | P | L | D | R | **PapR_7_ - A1R:dE_6_** |
| 0.588 ± 0.077 | **-** | F | dE | F | P | L | D | Y | **PapR_7_ - A1Y:dE_6_** |
| 1.264 ± 0.174 | **-** | F | dE | F | P | L | D | N | **PapR_7_ - A1N:dE_6_** |
| 0.213 ± 0.02 | **-** | F | dE | F | P | L | D | M | **PapR_7_ - A1M:dE_6_** |

IC_50_ values were calculated by GraphPad Prism 8, using the nonlinear inhibitor vs. normalized response method.

**Table S2:** Comparison between IC_50_ values of A1M-substituted known inhibitory peptides and their parent inhibitors, as determined by the lacZ-based reporter assay.

| IC_50_ [µM] | Peptide name |  |
| --- | --- | --- |
| 1.698 ± 0.111 | **E6A** | Position  **E6** |
| 0.485 ± 0.037 | **A1M:E6A** |  |
| 0.977 ± 0.04 | **dE_6_** |  |
| 0.213 ± 0.02 | **A1M:dE_6_** |  |
| 1.637 ± 0.102 | **F7A** | Position  **F7** |
| 0.482 ± 0.031 | **A1M:F7A** |  |
| 1.223 ± 0.07 | **dF_7_** |  |
| 0.58 ± 0.035 | **A1M:dF_7_** |  |

IC_50_ values were calculated by GraphPad Prism 8, using the nonlinear inhibitor vs. normalized response method.

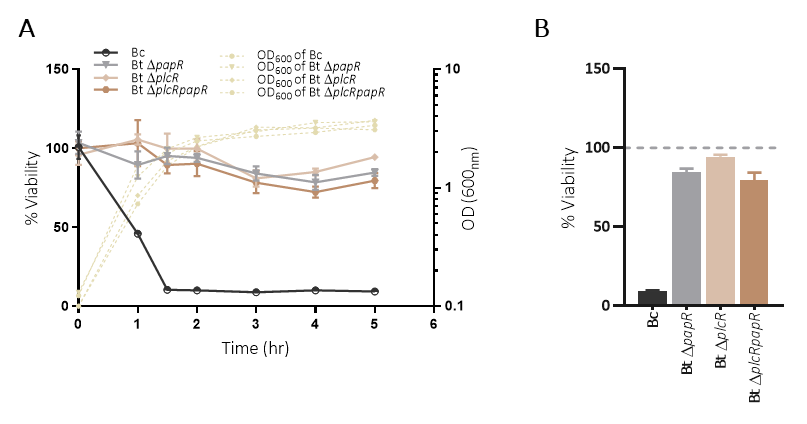

**Figure S2: PlcR regulon plays a major role in Bc** **cytotoxicity toward macrophages.** (A) Macrophage cells viability after challenge with culture supernatants from Bc*,* Bt ΔpapR A’Z, Bt ΔplcR and Bt ΔplcRpapR during different bacterial growth phase (data are representative of three independent experiments; mean ± SD, n = 3). (B) Macrophage cells viability after 5 hours represented in panel A (Data are representative of three independent experiments; mean ± SD, n = 3).

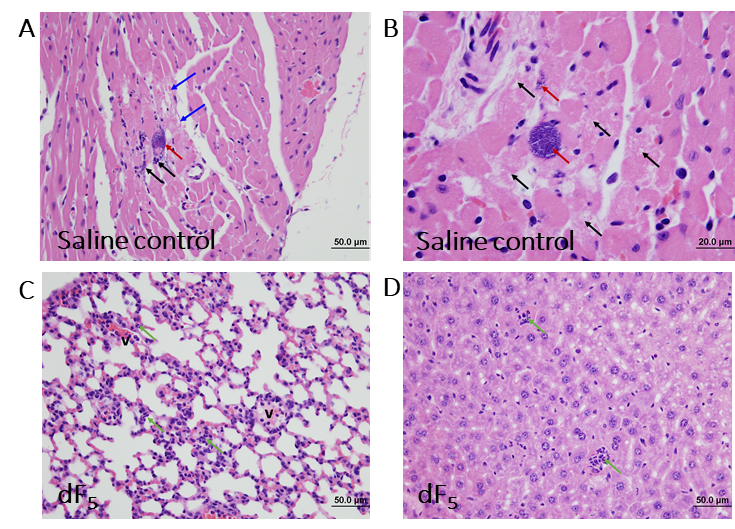

**Figure S3: Histopathological analysis of various tissues derived from mouse model of bacteremia caused by Bc** **strain, related to Figure 3**. (A-B) Representative images of histopathological changes in heart tissue of mice infected by Bc and treated with saline (H&E staining; A: 400x, B: 1000x; Black arrows, myocardial necrosis and cardiomyolysis. Red arrows, presence of bacterial colonies. Blue arrows: muscle fiber necrosis). (C-D) Representative images of histopathological changes in (C) lung and (D) liver tissue of mice infected by Bc and treated with PapR_7_–dF_5_ (H&E staining; C-D: 400x; Green arrows, infiltration of mildly increased neutrophils. V indicates blood vessel).

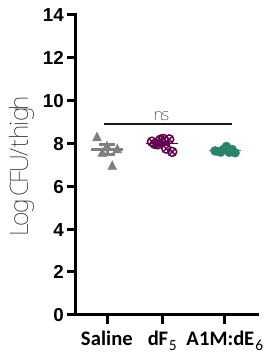

**Figure S4: PapR_7_ – A1M:dE_6_ relies on host defenses to clear bacteria during mouse thigh infection, related to Figure 4**. (A) Thigh bacterial load from neutropenic mice post-infection treated with saline, PapR_7_–dF_5_ and A1M:dE_6_ (mean ± SEM, n = 6-8). ns indicates no statistically significant difference between treatments.
